# Mutations in *Kinesin Family Member 6* Reveal Specific Role in Ependymal Cell Function and Human Neuro-Cranial Development

**DOI:** 10.1101/350140

**Authors:** Mia J. Konjikusic, Patra Yeetong, Rungnapa Ittiwut, Kanya Suphapeetiporn, John B. Wallingford, Christina A. Gurnett, Vorasuk Shotelersuk, Ryan S. Gray

## Abstract

Cerebrospinal fluid flow is crucial for neurodevelopment and homeostasis of the ventricular system of the brain, with localized flow being established by the polarized beating of the ependymal cell (EC) cilia. Here, we report a homozygous one base-pair deletion, c.1193delT (p.Leu398Glnfs*2), in the *Kinesin Family Member 6* (*KIF6*) gene in a child displaying neurocranial defects and intellectual disability. To test the pathogenicity of this novel human *KIF6* mutation we engineered an analogous C-terminal truncating mutation in mouse. These mutant mice display severe, postnatal-onset hydrocephalus. We generated a *Kif6-LacZ* transgenic mouse strain and report expression specifically and uniquely within the ependymal cell (EC) layer of the brain, without labeling other multiciliated mouse tissues. Analysis of *Kif6* mutant mice with scanning electron microscopy (SEM) and immunofluorescence (IF) revealed a reduction in EC cilia, without effect on other multiciliated tissues. Consistent with our findings in mice, defects of the ventricular system and EC cilia were observed in *kif6* mutant zebrafish. Overall, this work describes the first clinically-defined *KIF6* homozygous null mutation in human and defines KIF6 as a conserved mediator of neuro-cranial morphogenesis with a specific role in the maintenance of EC cilia in vertebrates.

**AUTHOR SUMMARY:** Cerebrospinal fluid flow is crucial for neurodevelopment and homeostasis of the ventricular system of the brain. Localized flows of cerebrospinal fluid throughout the ventricular system of the brain are established from the polarized beating of the ependymal cell (EC) cilia. Here, we identified a homozygous truncating mutation in *KIF6* in a child displaying neuro-cranial defects and intellectual disability. To test the function of KIF6 *in vivo*, we engineered mutations of *Kif6* in mouse. These *Kif6* mutant mice display severe hydrocephalus, coupled with a loss of EC cilia. Similarly, we observed hydrocephalus and a reduction in EC cilia in *kif6* mutant zebrafish. Overall, this work describes the first clinically-defined *KIF6* mutation in human, while our animal studies demonstrate the pathogenicity of mutations in *KIF6* and establish KIF6 as a conserved mediator of neuro-cranial development and EC cilia maintenance in vertebrates.

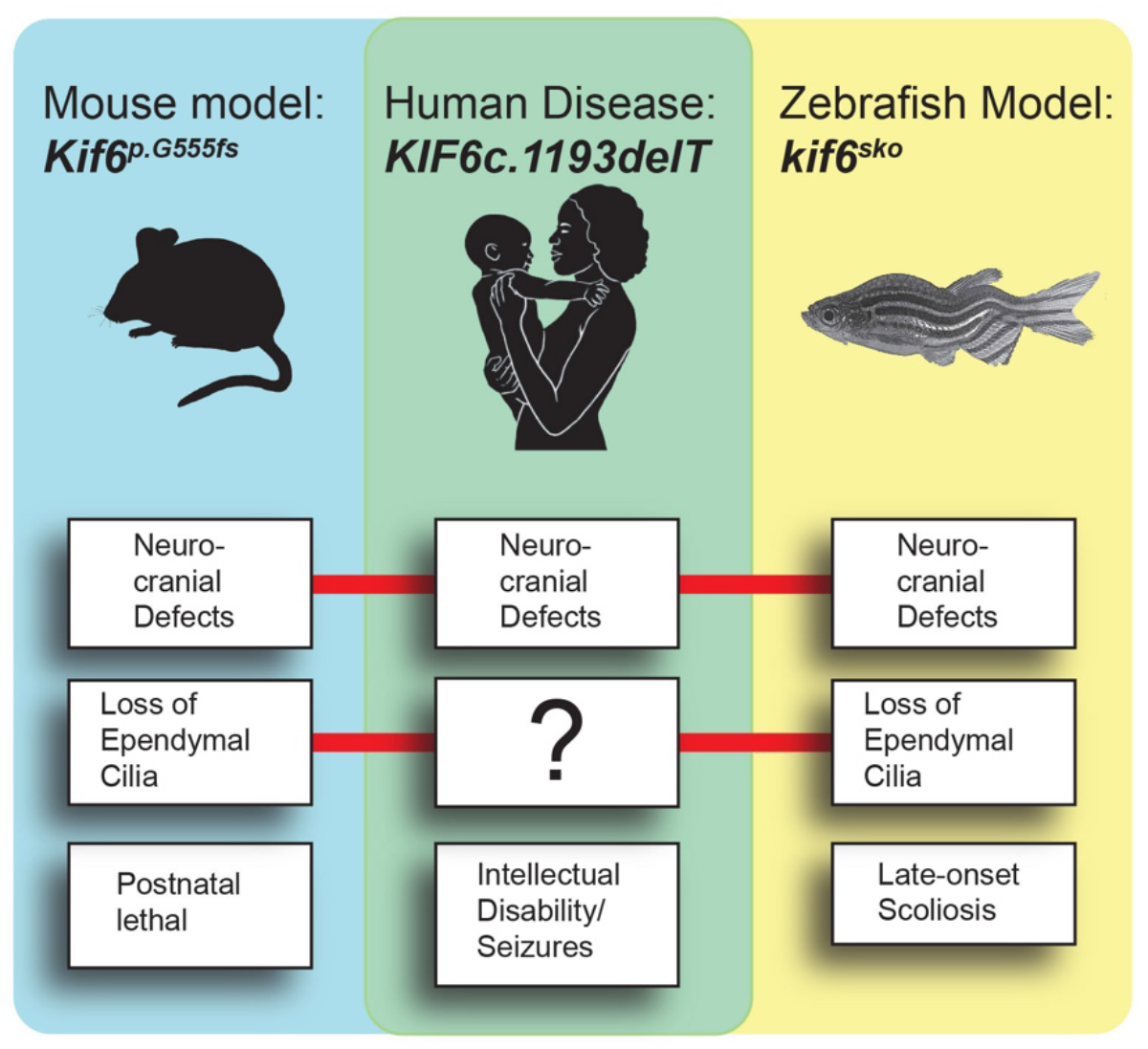

## INTRODUCTION

The delicate balance of cerebrospinal fluid (CSF) production and flow is important for the morphogenesis and function of the brain development and homeostasis. CSF circulation in human is largely due to gradients established by the secretion of CSF from the choroid plexuses, and its resorption at the arachnoid granulations [1]. The clinical significance of CSF stasis includes hydrocephalus and intracranial hypertension. Moreover, severely diminished CSF flow combined with increased intracranial pressure can secondarily cause ventriculomegaly, cognitive impairment, as well as degenerative and age-related dementias [2]. For these reasons, the identification of genetic risk factors involved in the pathogenesis of CSF stasis is critical for the development of genetic diagnostics and early interventions for these disorders.

One element for circulation of CSF is the multiciliated ependymal cells (ECs), which are specialized glial cells covering the ventricular walls of the brain and spinal canal [3]. In contrast, to primary cilia which are single, immotile cellular organelles extending from most cell types, ECs contain dozens of apically-arranged motile cilia, which beat in a polarized fashion to generate localized or near-wall CSF flows [4]. Defective EC cilia or loss of their polarized beating causes a disruption of this localized CSF flow leading to increased intracranial pressure, and dilation of ventricles and hydrocephalus in mice [5–8]. Importantly, this EC cilia-driven CSF flow is crucial for regulating brain function and adult neurogenesis [4, 9].

Impaired ciliary motility due to disruptions of the key kinesins, dyenins, and intraflagellar components necessary in most or all cilia, results in a syndromic condition known as primary ciliary dyskinesia (PCD) in humans [10, 11]. While hydrocephalus can occur in some PCD patients, it is a less common manifestation of the disease in humans [11]. In contrast, genes implicated in PCD or mutations which disrupt the structure or motility of all motile cilia are strongly correlated with hydrocephalus in mouse [8]. Alternatively, some hydrocephalus in mice with dysfunctional cilia may be the result of altered function of the choroid plexus, prior to the onset of cilia-driven CSF flow [7].

*KIF6* (Kinesin family member 6, MIM: 613919) encodes a member of the kinesin-9 superfamily of microtubules motor proteins which act predominately as “plus-end” directed molecular motors that generate force and movement across microtubules [12]. Kinesins are critical for numerous cellular functions such as intracellular transport and cell division, as well as for building and maintaining the cilium in a process known as intraflagellar transport [13]. During this process, kinesins have been shown to transport cargo within the ciliary axoneme [14], establish motility and compartmentalization of the axoneme [15], or to facilitate plus-end directed microtubule disassembly and control of axonemal length [16]. As such, multiple kinesins have shown to be associated with monogenic disorders affecting a wide-spectrum of tissues, with several modes of inheritance (www.omim.org). Interestingly, *KIF6* has previously been proposed as locus for susceptibility to coronary heart disease [17], while other studies did not substantiate this association [18]. We previously reported that *kif6* mutant zebrafish are adult viable exhibiting larval-onset scoliosis without obvious heart defects [19]. Because of these conflicting results, and a lack of relevant mouse models, the role of KIF6 in human disease remains an open question.

Here, we present a patient with consanguineous parents, presenting with neuro-cranial defects and intellectual disability. Homozygosity mapping followed by whole-exome sequencing (WES) identified a novel homozygous frameshift mutation in *KIF6* which is predicted to result in the truncation of the C-terminal cargo-binding domain of the kinesin motor protein. We generated an analogous frameshift mutation in the mouse and found that these mutant mice displayed progressive, postnatal-onset hydrocephalus with cranial expansion, coupled with a reduction in the quantity of EC cilia. In addition, we observed that *kif6* mutant zebrafish also display dilation of the ventricular system, coupled with reduced EC cilia. Interestingly, we failed to observe additional PCD related defects of other multiciliated tissues in *Kif6* mutant mouse or zebrafish models. Together these results demonstrate that KIF6 function is unique and specific for EC cilia. Finally, we propose that *KIF6* represents a novel gene for neuro-cranial development and intellectual disability in humans.

## RESULTS

### Clinical features and mutation identification

We identified a Thai boy with intellectual disability and megalencephaly. His parents were first cousins. He was born at 34 weeks gestation with a head circumference of 34 cm (97^th^ centile). APGAR scores were 7 and 9 at 1 and 5 minutes, respectively. Neonatal hypoglycemia (blood sugar of 11 mg/dL) and neonatal jaundice were treated promptly. In the first few months of life, he was found to have delayed neurodevelopment and central hypotonia. He was able to hold his head at 5 months, rolled over at 8 months, walked and had first words at 2 years old. At the age of 9 years and 9 months, an IQ test by Wechsler Intelligence Scale for Children: 4th edition (WISC-IV) revealed that his full-scale IQ was 56, indicating intellectual disability. The patient had possible seizure activity at age 10 described as parasomnias, was found to have intermittent bifrontocentreal rhythmic theta activity, and the spells resolved after valproic acid therapy. His height and weight followed the curve of 50th centile, but his head circumference remained at 97th centile (53.5 cm and 55 cm at 6 and 9 years old, respectively). Physical examination was generally unremarkable except macrocephaly and low-set prominent anti-helical pinnae (Fig 1A). Eye examination, hearing tests, thyroid function tests, chromosomal analysis, and nerve conduction velocity were normal. Both brain CT scans at 4 months and 8 years old and brain MRI at 7 months old showed a slight dolichocephalic cranial shape (cephalic index = 75), without overt structural brain abnormalities (Fig 1B-D). X-ray analysis of the spine showed no obvious scoliosis at 10-years-old (Fig 1E).

**Fig 1.**
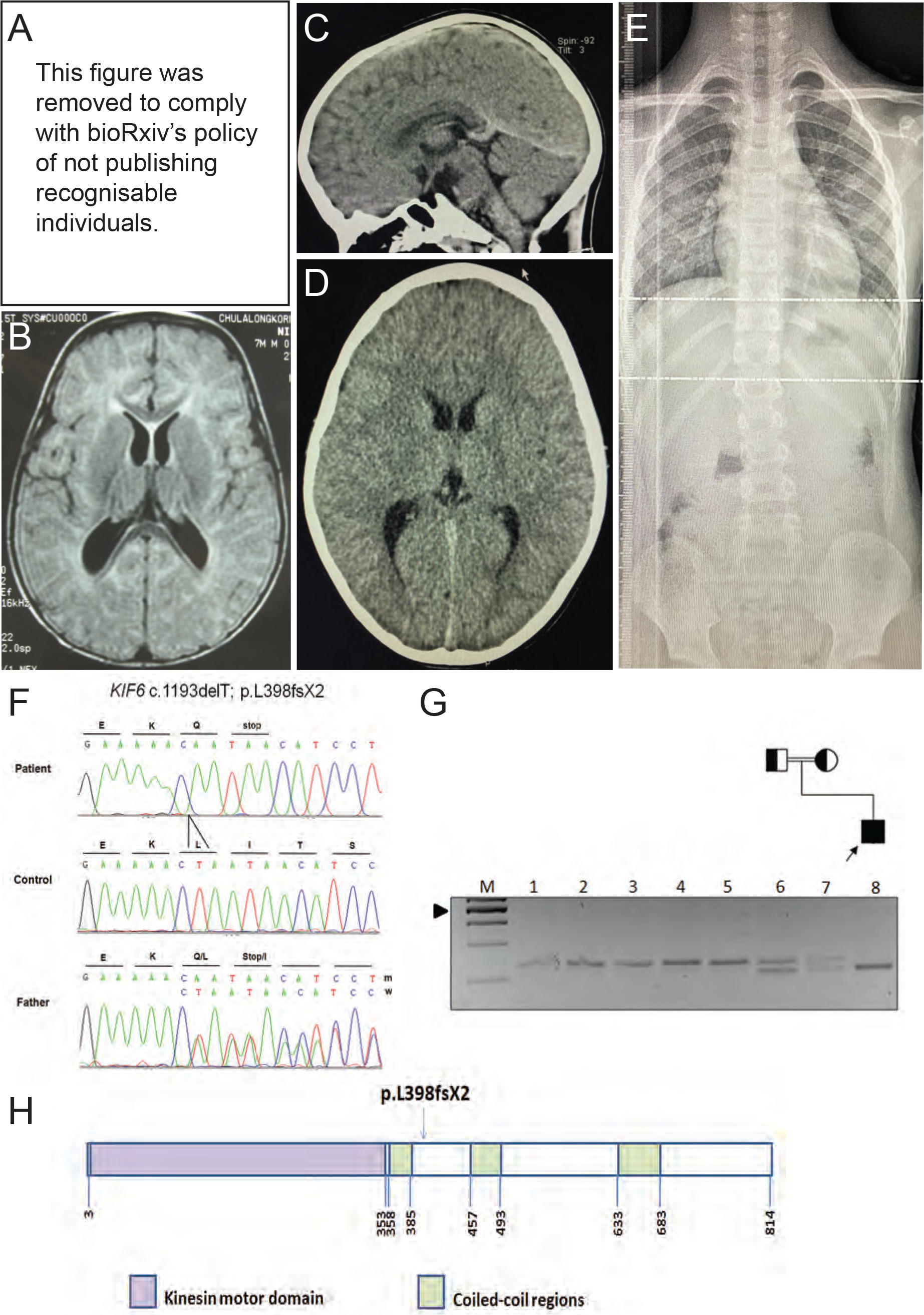
*KIF6* mutation in a child with intellectual disability. (A) A low-set prominent antihelical left pinna. (B) MRI of the brain at 7 months old shows dolichocephaly with a normal brain structure. (C) and (D) CT of the brain at 8 years old, sagittal and axial views, respectively show dolichocephalic shape of the cranium (cephalic index =75) without demonstrable intracranial abnormality. (E) X-ray of the spine shows no scoliosis (F) Electropherograms of the patient, a control, and the patient’s father in the upper, middle and lower panels, respectively. The patient is homozygous while his father is heterozygous for the c.1193delT. (G) Pedigree and RFLP, using MfeI restriction enzyme: Lane M = 100 bp marker. The arrow head indicates the 500 bp band. Lanes 1-5 are controls. Lanes 6 and 7 are the proband’s father and mother, respectively, showing that they are heterozygous. Lane 8 is the proband showing that he is homozygous for the c.1193delT. (H) Representative KIF6 structure. The arrow shows the position of the c.1193delT mutation.

To elucidate the genetic etiology, we performed homozygosity mapping, whole genome array comparative genomic hybridization (CGH), and whole exome sequencing (WES). WES identified 83 homozygous variants, which had not been reported as SNPs in dbSNP137 (S1 Table). We then selected only those located within the 63 homozygous regions found by homozygosity mapping (S2 Table). Seven candidate variants (one frameshift and six missense mutations; Table I) were identified. Of the six missense, five were predicted to be either benign by Polyphen-2 or tolerated by SIFT prediction programs. The remaining variant, c.235G>A; p.V79M of the *Carboxypeptidase E* (*CPE*) gene, was not evolutionarily conserved among diverged species (S1 Fig). We, therefore, decided to further our study on the only candidate truncating mutation, a homozygous one base-pair deletion, c.1193delT (p.Leu398Glnfs*2) in exon 11 of *Kinesin family member 6* (*KIF6*) (NM_001289021.2).

*KIF6* is located on human chromosome 6p21.2 and comprises 23 exons. The 2.4-kb *KIF6* cDNA encodes a canonical N-terminal kinesin motor domain (amino acid positions 3-353) and three coiled-coil regions (amino acid positions 358-385, 457-493, and 633-683), predicted by SMART server [20]. Segregation of the homozygous sequence variant with the disease phenotype was confirmed by Sanger sequencing (Fig 1F) and by restriction fragment length polymorphism (RFLP) analysis of the pedigree (Fig 1G), while his parents and his unaffected brother were heterozygous for the deletion (Fig 1F, G and data not shown). The deletion was not observed in our 1,600 in-house Thai exomes, the 1000 Genome Database, and the ExAC Database. The pedigree combined with the novelty of the mutation in *KIF6* presented here, strongly suggest this C-terminal truncating mutation in KIF6 may be etiologic for neuro-cranial developmental defects.

**Table 1.** Seven candidate variants from WES and homozygosity mapping.

| # | Chromosome<br>(position) | Gene | Zygoty | Nucleotide<br>change | Amino acid change | Polyphen-2<br>(score) | SIFT |
| --- | --- | --- | --- | --- | --- | --- | --- |
| 1 | 6 (39513453) | *KIF6* | Homozygous | c.1193delT | p.L398QfsX399 |  |  |
| 2 | 4 (166300608) | *CPE* | Homozygous | c.235G>A | p.V79M | Possibly<br>damaging<br>(0.889) | Deleterious |
| 3 | 6 (41895234) | *BYSL* | Homozygous | c.391C>T | p.R131C | Probably<br>damaging<br>(0.975) | Tolerated |
| 4 | 7 (44120345) | *POLM* | Homozygous | c.359G>A | p.R120Q | Probably<br>damaging<br>(0.999) | Tolerated |
| 5 | 7 (64167644) | *ZNF107* | Homozygous | c.962T>G | p.I321S | Probably<br>damaging<br>(0.969) | Tolerated |
| 6 | 2 (96148317) | *TRIM43B* | Homozygous | c.146C>T | p.P49L | Benign<br>(0.071) | Tolerated |
| 7 | 5 (140307142) | *PCDHAC1* | Homozygous | c.665T>C | p.I222T | Benign<br>(0.001) | Tolerated |

### Generation of *Kif6* Mutation in Mouse

To test the functional consequence of the C-terminal truncating p.L398fsX2 mutation (Fig 1H), we generated an analogous frameshift mutation in exon14 of the mouse *Kif6* (ENSMUST00000162854) gene, which is ~150bp downstream of the frameshift mutation found in the patient (Fig 2A). After backcross of founder mice to C57B6/J strain, we identified a nonsense allele with scarless insertion (c.1665ins) of a 3-stop donor cassette-providing integration of an ochre termination codon in all three reading frames into the endogenous *Kif6* locus (S2 Fig). This endonuclease-mediated insertional frameshift mutation (*Kif6*^*em1Rgray*^) is predicted to truncate the C-terminal cargo-binding domain of the kinesin motor protein (p.G555+6fs). This novel mutant allele of *Kif6* (hereafter called *Kif6*^*p.G555fs*^) is predicted to encode a C-terminal truncated KIF6 protein 168 amino acids longer than is predicted for the human p.L398fsX2 variant (Fig 2A).

**Fig 2.**
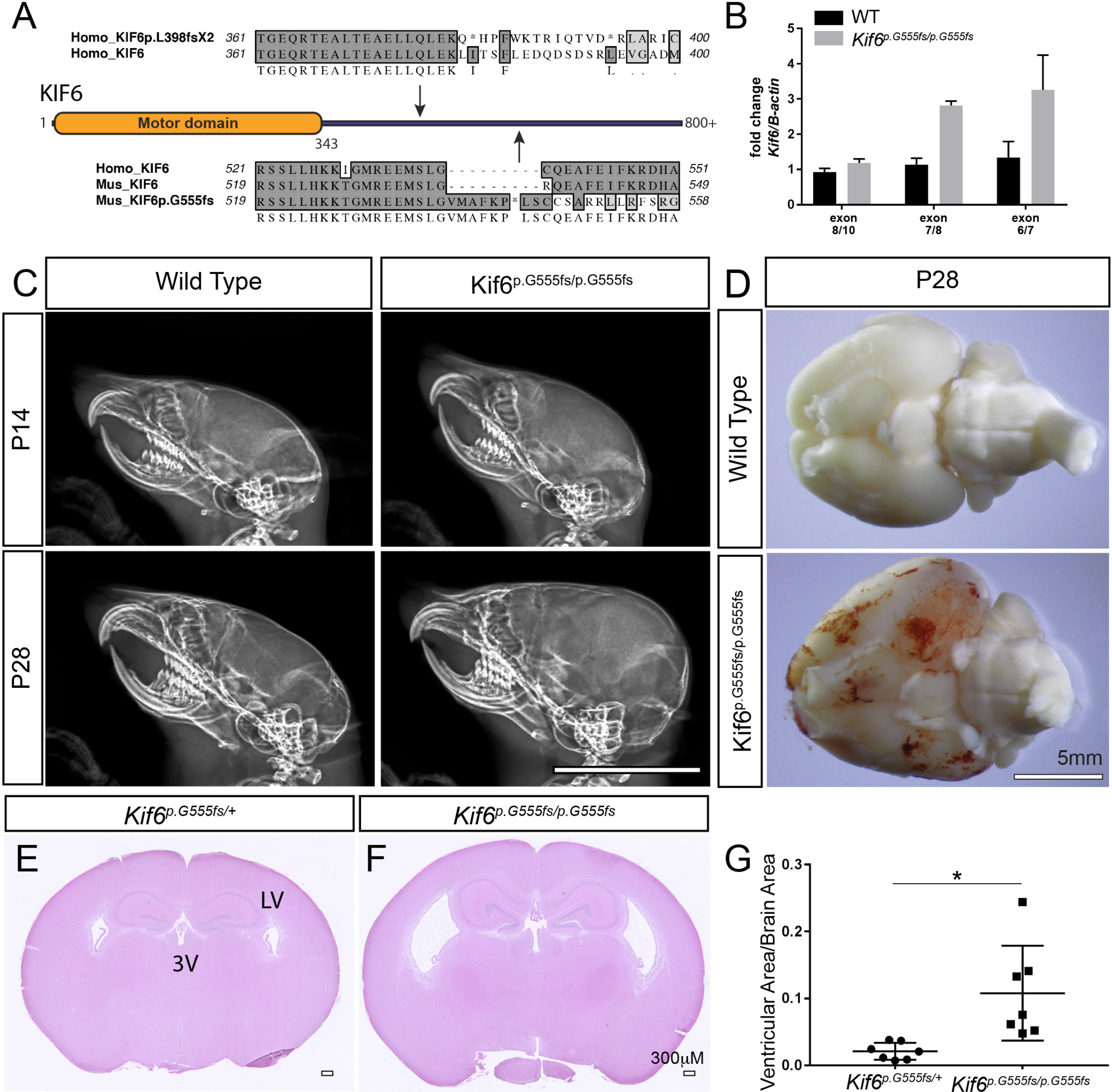
*Kif6*^*p.G555fs*^ mutant mice display progressive hydrocephaly. (A) Schematic of the nonsense mutation in the patient (*KIF6*^*p.L398fsX2*^) and the mouse mutation (*Kif6*^*p.G555fs*^), both predicted to truncate the C-terminal domain of KIF6 protein. (B) Fold change qRT-PCR of *Kif6* expression using cDNA libraries derived from lateral ventricles from WT (black bars) and *Kiffl*^*p.G555fs/p.G555fs*^ (gray bars) mutant mice. (C) Lateral X-rays of mouse cranium at P14 and P28 showing the progressive cranial expansion in *Kif6*^*p.G55/pG555fs*^ homozygous mutant mice. (D) Ventral view of whole mouse P28 brain to highlight hemorrhaging and slight enlargement of total brain size in *Kif6*^*p.G55/p.G555fs*^ homozygous mutant mice. (E, F) H&E stained coronal sections of the mouse brain (P14), showing dilation of the lateral (LV) and third (3 V) ventricles in *Kif6*^*p.G55/p.G555fs*^ homozygous mutant mice (F). (G) Quantitation of ventricular area over total brain area in *Kif6*^*p.G55/p.G555fs*^ homozygous mutant mice and heterozygous littermate controls (n=7 mice per genotype; two-tailed t-test; p=0.0173). Scale bars: 1cm in (C); 5mm in (D); and 300 μin (F,F).

### Hydrocephalus in *Kif6*^*p.G555fs*^ Mouse

Intercrossing *Kif6*^*p.G5555s/+*^ heterozygous animals gave offspring with the expected Mendelian ratios, with typical appearance at birth. However, beginning at postnatal day (P)14-onwards, 100% (n=7) of *Kif6*^*p.G555fs/p.G555fs*^ homozygous mutant mice displayed classic indications of hydrocephalus including doming of the cranium (Fig 2C), a hunched appearance, and with decreased open field activity. We observed apparent megalencephaly and hemorrhaging in older (P21-P28) *Kif6*^*p.G555fs/p.G555fs*^ mutant brains (Fig 2D), which likely results from increased intracranial pressure and swelling of the ventricles causing damage to the neural tissue against the cranium. At P14, the body weights were not significantly decreased in *Kif6*^*p.G555fs*^ mutants (5.8±1.3 (g)rams) compared with littermate controls (7.0±1.2g) (n=5/genotype; *p*=0.17). However, at P28 mutant mice showed decreased weight on average (12.67±1.53 g) compared to littermate controls (15.33±1.15g), although this trend was not statistically significant (n=3/genotype; *p*= 0.07). At P28, extracted whole brain sizes appear to be larger in *Kif6*^*p.G555fs/p.G555fs*^ mutants compared to non-mutant littermate controls (Fig 2D). For these reasons, mutant animals were not maintained for observation past P28. qPT-PCR analysis of several *Kif6* exon-exon boundaries found no evidence for non-sense mediated decay in *Kif6*^*p.G555fs*^ mutant mice (Fig 2B).

To determine whether a more N-terminal truncated *Kif6* mutation would result in a more severe hydrocephalus phenotype, we isolated a conditional-ready *Kif6* allele, where exon 4 is flanked by LoxP sites (*Kif6*^*tm1c*^) (KOMP repository, see Methods and Materials). Recombination of the *Kif6*^*tm1c*^ allele is predicted to generate a frameshift mutation, which should generate a severely truncated, 89 amino acid, KIF6 protein (p.G83E+6fs) with a non-functional N-terminal motor domain. We generated a whole body conditional knockout by crossing the *Kif6*^*tm1c*^ mouse to the *CMV-Cre* deleter mouse [21]. We observed postnatal-onset, hydrocephalus in *CMV-Cre; Kif6*^*tm1c/tm1c*^ conditional mutant mice (n=10) analogous to our observations in *Kif6*^*p.G555fs/p.G555fs*^ mutant mice (data not shown). Interestingly, we find no evidence of non-sense mediated decay in these mutant mice despite the generation of an early premature termination codon (data not shown). Because the onset and progression of hydrocephalus was equivalent comparing the whole-body conditional *CMV-Cre; Kif6*^*tm1c/tm1c*^ and *Kif6*^*p.G555fs/p.G555fs*^ mutant mice strains we suggest that any KIF6 protein encoded by these mutant mouse strains is likely non-functional. Given its relevance to the human mutation, the majority of experiments were all done using the *p.G555fs* allele.

Mouse brains were analyzed histologically by hematoxylin and eosin (H&E) stained coronal sections. Our analysis of coronal sectioned brain at P14 failed to find significance when comparing the total area in section (499.2+39.9μM (Control) vs. 552.5+50.8μM (*Kif6*^*p.G555fs/p.G555fs*^); n=7/genotype; *p*=0.42). However, lateral and third ventricles (LV and 3V respectively) were obviously enlarged in *Kif6*^*p.G555fs/p.G555fs*^ mutants (Fig 2F). Quantitation of LV volumes normalized to total brain volume confirmed ventricular expansion in *Kif6*^*p.G555fs/p.G555fs*^ mutants (n=7 for each genotype; p≤0.05; Fig 2G). No obvious defects of the cortex or development of other brain regions in *Kif6*^*p.G555fs/p.G555fs*^ mutant mice were apparent at a gross anatomical level (Fig 2E, F). Together these data suggest that *Kif6*^*p.G555fs/p.G555fs*^ mutant mice display postnatal-onset, progressive hydrocephalus, without obvious overgrowth of neural cortex.

### *Kif6* is Expressed Specifically in the ECs of the Mouse Brain

To determine the endogenous expression patterns of *Kif6* in the mouse, we isolated a *Kif6-LacZ* reporter mouse (*Kif6-LacZ*^*tm1b*^) (KOMP repository, see Methods and Materials). Hemizygous *Kif6-LacZ*^*tm1b/+*^ mice appeared unremarkable and exhibited no evidence of hydrocephalus. Intercrosses of *Kif6*^*tm1b/+*^ hemizygous mice failed to generate litters with *Kif6*^*tm1b/tm1b*^ homozygous mice, suggesting that the homozygosity of the *lacZ* expressing allele is embryonic lethal (data not shown). At P10 and P21, *Kif6*^*tm1b/+*^ transgenic mice showed *lacZ* expression throughout the ependyma of the ventricular system including the central canal. However, no *lacZ* expression was detected in the choroid plexus or in other regions of the brain (Fig 3A, A’ and S3B’ Fig), with the exception of a small population of cells flanking the third ventricle (arrows, S3 Fig). Interestingly at P10, other multi-ciliated tissues in these transgenic mice such as the oviduct or trachea were not labeled (Fig 3B-C’). Moreover, no laterality defects or obvious changes to trachea cilia were observed in *Kif6*^*p.G555fs/p.G555fs*^ mutant mice (S6A, B Fig), suggesting that *Kif6* expression and function are tightly restricted to the multiciliated EC in mouse. Taken together these data suggested a cellular mechanism centered on defective ECs underlying the development of hydrocephalus in *Kif6*^*p.G555fs/p.G555fs*^ mutant mice.

**Fig 3.**
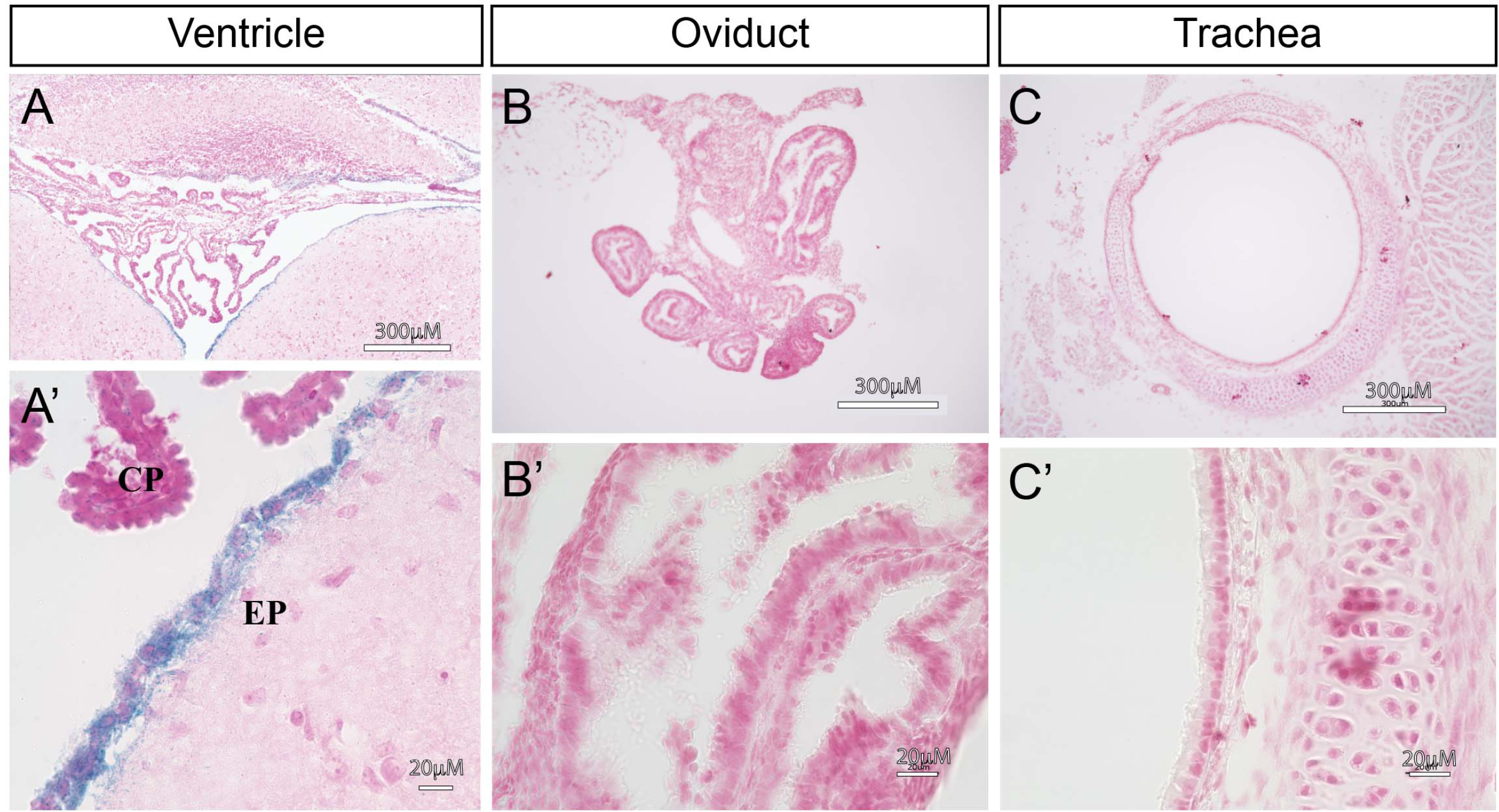
*Kif6-LacZ* expression is specific to the ependymal cells. (A-C’) Representative LacZ staining in a variety of multiciliated tissues from P10 *Kif6-LacZ*^*tm1b/+*^ transgenic mice. (A, A’) Coronal section at the 4^th^ ventricle showing specific *LacZ* expression in the ependymal cell (EC) layer and stark lack of expression in the choroid plexus (CP) or surrounding neuronal tissues. (B, B’) Sectioned oviduct tissue shows no *LacZ* expression. (C, C’) Sectioned trachea tissue shows no *LacZ* expression. Scale bars: 300μM in (A-C); and 20μM in (A’-C’).

### Progressive Loss of Cilia in *Kif6*^*p.G555fs/p.G555fs*^ Mutant Mice

Defects in ECs and their cilia are known to cause hydrocephalus in mouse [8]. To assay EC cilia, we utilized scanning electron microscopy (SEM) to directly visualize the LV. Heterozygous *Kif6*^*p.G555fs/+*^ mice displayed a high-density of regularly spaced EC multiciliated tufts along the LV surface (Fig 4A-A’), typical for P21 mice [22]. In contrast, homozygous *Kif6*^*p.G555fs/p.G555fs*^ mutant mice displayed a marked reduction of multiciliated tufts across the LV wall, coupled with a reduction in the density of ciliary axonemes extending from the ECs (Fig 3B-B’). The loss of EC cilia was more severe at P28 (S4 Fig). Together, these data suggested that hydrocephalus may result from either a reduction in EC differentiation and/or defects in EC cilia maintenance during postnatal development.

**Fig 4.**
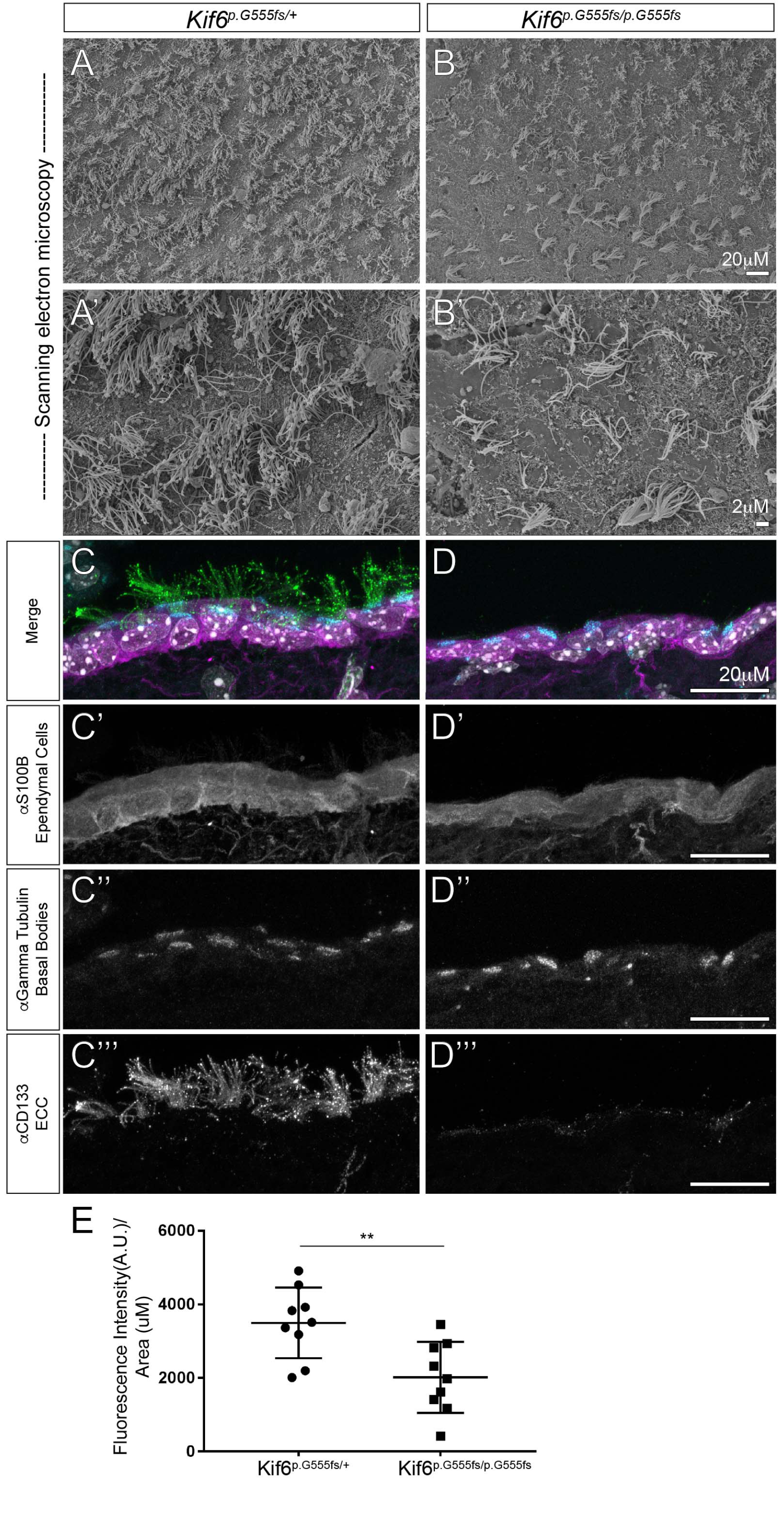
*Kif6* mutant mice have a reduction in ependymal cell cilia. (A-B’) SEM of the lateral ventricular wall (*en face* view) in *Kif6*^*p.G55/pG555fs*^ homozygous mutant mice and heterozygous littermate controls at P21, showing a reduction in the number and density of EC cilia tufts. (C-D”’) Immunofluorescence of the wild-type (WT) and *Kif6*^*p.G55/p.G555fs*^ homozygous mutant mice at P21. (C-D) Three color merge of (C’, D’) αS100B (ependymal cells; magenta) channel; (C”, D”) α-γ-tubulin (basal bodies; cyan) channel; and (C”’, D”’) αCD133 (prominin-1; green). (C’, D’) αS100B staining showing no changes in ependymal cell specification between *Kif6*^*p.G55/p.G555fs*^ homozygous mutant and WT mice. (C”, D”) α-γ-tubulin staining showing typical basal body positioning at the apical surface of ECs in both *Kif6*^*p.G55/p.G555fs*^ homozygous mutant and WT mice. (C”’, D”’) αCD133 staining reveals a marked of EC cilia projecting into the ventricular lumen in *Kif6*^*p.G55/p.G555fs*^ homozygous mutant mice compared to WT mice. (E) Quantitation of fluorescent intensity of the CD133 channel (ciliary axonemes) as the average line scan value along the lumal surface of mouse ventricles. Scale bars: 20μM in (A, B); 2 μM in (A’, B’); and 20 μM in (C-D”’).

To address the differentiation status of the ECs, we utilized immunofluorescence (IF) in coronal sectioned brain tissues to image known proteins components of the EC and their cilia. At P21, we observed the expression of the ependymal cell-marker S100B [22] throughout the epithelium lining luminal surface of the ventricles, as well as, the presence of apically localized g-tubulin-positive basal bodies within these ECs in both WT (Fig 4C-C”) and *Kif6*^*p.G555fs/p.G555fs*^ mutant mice (Fig 4D-D”). Conversely, we observed a severe reduction in the density of CD133-positive EC axonemes [23] extending into the ventricular lumen in *Kif6^p.G555fs/p.G555fs^* mutant mice (Fig 4C, C”’), compared with WT (Fig 4D, D”’). Quantitation of several sections from independent mice confirmed a severe reduction of CD133-positive ciliary axonemes due to the loss of KIF6 function (n=5 mice/genotype, p<0.001) (Fig 4E). These results suggest that the onset of hydrocephalus in *Kif6* mutant mice is primarily due to the loss of EC ciliary axonemes and not the result of defects in the differentiation of these cells.

### Ventricular Dilation and Reduced EC Cilia in *kif6* Mutant Zebrafish

Previous studies in *kif6*^*sko/sko*^ mutant zebrafish found late-onset scoliosis, without obvious hydrocephalus or defects in EC cilia during early embryonic development [19]. Interestingly, reduced CSF flow, ventricular dilation, and loss of EC cilia during larval zebrafish development is associated with scoliosis [24]. In order to determine if *kif6* mutant zebrafish display changes in the ventricular system later in adults, we used iodine contrast-enhanced, micro computed tomography (μCT) [25] to generate high-resolution (5 μM) images of the intact zebrafish brain. After reconstruction and alignment of 3D tomographic datasets in the coronal plane, we utilized stereotyped landmarks of the zebrafish brain and spinal cord [26] to compare equivalent axial sections of aged matched (90 days post fertilization (dpf)) *kif6*^*sko/+*^ heterozygous and *kif6*^*sko/sko*^ homozygous mutant zebrafish. At each axial level of the brain (Fig 5A), we observed consistent dilation of the ventricular system and central canal in *kif6*^*sko/sko*^ homozygous mutant zebrafish (yellow arrows; Fig 5C, E, G), compared to a stereotyped anatomy of the wild-type (WT) zebrafish brain (Fig 5B, D, F). Multiple regions of *kif6*^*sko/sko*^ mutant zebrafish brain were found to be structurally abnormal in *kif6*^*sko/sko*^ mutants compared to WT zebrafish (S1, 2 Movies). We next quantified the areas of two anatomically distinctive ventricles in our tomographic datasets: (i) the tectal ventricle (TecV) and (ii) a region of the rhombencephalic ventricle (RV) just posterior to the lobus facialis [26]. We found that both the TecV and the posterior RV were significantly more dilated in *kif6*^*sko/sko*^ mutant zebrafish comparing several optical sections from independent aged-matched zebrafish (n=3 fish/genotype; p<0.0001). The central canal was also clearly dilated in *kif6*^*sko/sko*^ mutant zebrafish (yellow arrow, Fig 5G). However, we were unable to reliably quantify this area in WT samples at the current resolution. Our previous observations in *kif6*^*sko/sko*^ mutant zebrafish embryos failed to find phenotypes that are characteristic of cilia defects, such as hydrocephalus, *situs inversus*, or kidney cysts [19]. Moreover, we observed normal development and function of EC cilia in the central canal in embryonic mutant zebrafish [19]. These data, together with our new observations of ventricular dilation in adult *kif6* mutants (Fig 5), suggest that Kif6 is required for the post-embryonic maintenance of the EC cilia as was observed in other zebrafish mutants displaying similar scoliosis as observed in *kif6* mutant zebrafish [19, 24].

**Fig 5.**
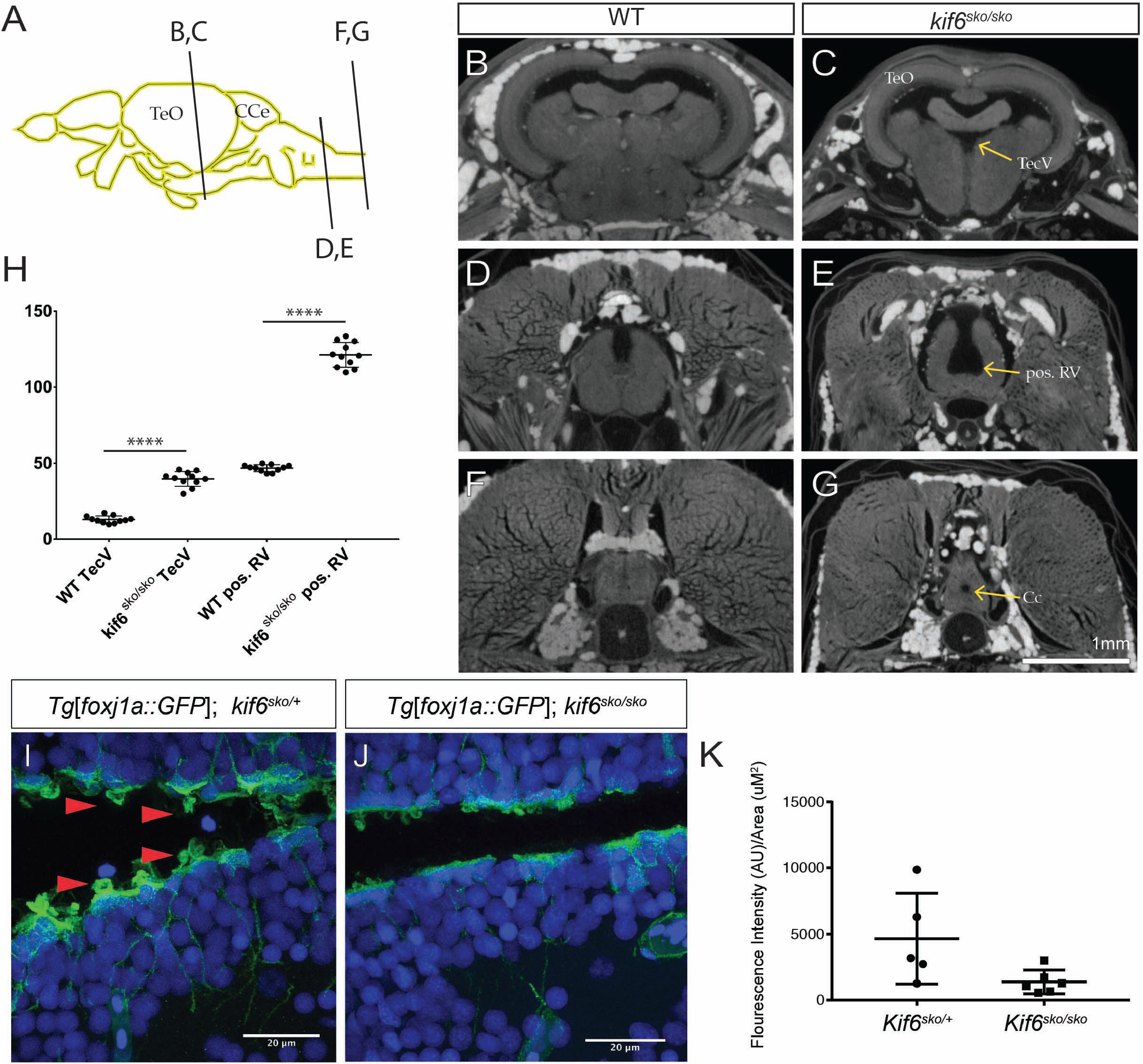
*kif6* mutant zebrafish display dilation of the ventricular system and loss of ependymal cell cilia. (A) Schematic of adult zebrafish brain highlighting the relative sectioning of the zebrafish brain. (B-G) Coronally aligned, optical sectioning of Iodine-contrasted μCT imaging datasets of WT (B, D, F) and *kif6*^*sko/sko*^ homozygous mutant (C, E, G) zebrafish brain at 90dpf. (B-C) The medial region of the TeO showing the TecV which is dilated in *kif6*^*sko/sko*^ mutant fish (C) compared to age-matched WT (B). (D-E) Sectioning at the region of the medulla oblongata posterior to the lobus facialis showing dysmorphogenesis and deepening of the RV in *kif6*^*sko/sko*^ mutants (E) compared with WT (D) zebrafish. (F-G) Spinal cord sectioning showing dilation of the Cc in *kif6*^*sko/sko*^ mutant (G) compared to WT (F) zebrafish. Scale Bars: 1mm in (B-G and 20 μM in (I, J). (H) Quantitation of the area of the TecV and the RV posterior to the lobus facialis (pos. RV) in WT and *kif6*^*sko/sko*^ mutant zebrafish, highlighting the consistent dilation in *kif6*^*sko/sko*^ mutants (n=11 sections/genotype; two-tailed t-test; ****, p<0.0001). (I-J) Immunofluorescence of *Tg[foxj1a::GFP]*in both heterozygous and homozyogus *kif6*^*sko/sko*^ mutant zebrafish stained with αGFP (green) and DAPI (blue) showing GFP positive ECs in both genotypes. However, *kif6^sko/^+* heterozygous zebrafish display numerous apical tufts of cilia (red arrows) projecting into the ventricle lumen, which are markedly reduced in *kif6*^*sko/sko*^ mutant zebrafish. Scale Bars: 1mm in (B-G); and 20 μM in (I, J). *TecV-tectal ventricle TeO-tectum opticum; CCe-corpus cerebelli; RV-rhombencephalic ventricle ventricle; and Cc-central canal.*

In order to assay whether EC cilia were affected in adult (90dpf) *kif6*^*sko/sko*^ mutant zebrafish, we isolated a stable transgenic allele, *Tg(Foxj1a:GFP)*^*dp1*^ which effectively labels multiple multiciliated *Foxj1a*-positive cell lineages, including ECs, with cytoplasmic EGFP in zebrafish [24]. We observed no differences in the specification of *Foxj1a:GFP*-positive ECs comparing WT and homozygous *kif6*^*sko/sko*^ mutant fish (Fig 5I, J). Cytoplasmic GFP can freely diffuse into and label the ciliary axoneme [27]. As such, we were able to observe GFP-positive EC cilia projecting into the ventricular lumen in *Tg(Foxj1a:GFP)*^*dp1*^; *kif6*^*sko/+*^ heterozygous fish (red arrowheads; Fig 5I). In contrast, these GFP-labeled EC cilia were reduced or absent in *Tg[Foxj1a:GFP]*^*dp1*^; *kif6*^*sko/sko*^ mutant fish (Fig 5J, K). Furthermore, SEM analysis of the ventricles in *kif6*^*sko/sko*^ mutant zebrafish further supported our observations of ventricular dilation and loss of EC cilia in adult *kif6*^*sko/sko*^ mutant fish (S5 Fig). Akin to our observations in *Kif6* mutant mice trachea, we did not observe defects of other multiciliated tissues such as the nasal cilia (S6C, D Fig) in *kif6*^*sko/sko*^ mutant zebrafish. Together, these data suggest that Kif6 functions specifically in the maintenance of EC cilia as well as for ventricular homeostasis in zebrafish.

## DISCUSSION

This study demonstrates the importance of *KIF6* for neuro-cranial development in vertebrates, and a unique and highly specialized role in ependymal cells where its function is important for maintenance of EC cilia. This is supported by several lines of evidence including the discovery of a novel nonsense-mutation of *KIF6* in a child with neuro-cranial defects and intellectual disability and underscored by functional analysis in both mouse and zebrafish *Kif6* mutant models (Table 2).

**Table 2.** KIF6 mutations discussed in this paper.

| ID/Allele | Species | Nucleotide change | Amino acid change | Phenotype reported | Reference |
| --- | --- | --- | --- | --- | --- |
| Patient | Human | c.1193delT | p.L398QfsX399 | Delayed neurodevelopment and central hypotonia, neuro-cranial defects, and intellectual disability. | this work |
| *Kif6*<sup>p.G555fs/p.G555fs</sup> | Mouse | c.1665ins | p.G555+6fs | Severe progressive hydrocephalus, loss of ependymal cell cilia | this work |
|  | Mouse | deletion of exon 4 | p.E83+39fs | Severe progressive hydrocephalus, loss of ependymal cell cilia | this work |
| *Kif6*<sup>Δ3/Δ3</sup> | Mouse | c.177-251_del | p.Lys59Asn, Phe60_Ser84_del | None reported | [28] |
| *kif6*<sup>sko/sko</sup> | Zebrafish | c.205C>A | p.Tyr53X | Larval onset scoliosis, hydrocephalus, loss of ependymal cilia | [19], this work |

We identified a homozygous *KIF6* c.1193delT mutation in a child with macrocephaly and cognitive impairment that segregated with this phenotype in his family, and leads to a loss of the C-terminal second and third coiled-coil regions which are important for dimerization and cargo selectivity of kinesin motors [13]. We engineered an analogous, C-terminal truncating mutation of *KIF6* in mouse, which displays severe hydrocephalus and defects of EC cilia providing strong evidence for pathogenicity of the mutation in the child. Other than the case described here, no prior mutation directly attributed to human disease has been described for *KIF6*. Taken together, the clinical data reported here suggest that biallelic mutations in *KIF6* may underlie some unexplained intellectual disability and neuro-cranial developmental defects. Future analyses of *KIF6* mutations in these patient groups are warranted.

Further, our analysis of several independent loss-of-function *Kif6* mutant animal models found no evidence of obvious heart abnormalities to explain the prior association of the common variant *KIF6* p.W719R in some[17], but not all [18], studies of coronary heart disease in humans. Because expressed sequence tag clones of *KIF6* have not been reported from cDNAs libraries derived from human heart or vascular tissues (UniGene 1956991 - Hs.588202), any possible functional effects of KIF6 on heart function remains unexplained. However, detailed analysis of coronary function was not explored in our models, therefore it is possible that subtle defects may be present.

An ENU-derived *Kif6* splice acceptor site mutant mouse strain, predicted to delete the 3rd exon of *KIF6* (*Kif6*^*Δ3/Δ3*^), also did not show cardiac or lipid abnormalities [28]. Of note this mutant mouse was also not reported to have hydrocephalus. Our analysis shows that the loss of exon 3 in *Kif6*^*Δ3/Δ3*^ mutant mouse generates an inframe deletion of only 25 amino acids in the N-terminal motor domain of the KIF6 protein, otherwise generating a mostly full-length KIF6 protein (Table 2). In contrast, here we report two novel *Kif6* mutant mice: (i) a C-terminal *Kif6*^*p.G555fs/p.G555fs*^ deletion mutant, predicted to truncate 248 amino acids of the C-terminal domain, which are important for cargo binding in Kinesin motor proteins [13]; and (ii) a conditional *CMV-Cre;Kif6*^*tm1c/tm1c*^ mutant which recombines exon 4 leading to an early frame shift mutation predicted to generate a N-terminal truncated 122 amino acid KIF6 protein (Table 2), both of which display indistinguishable progressive, hydrocephalus. The most parsimonious explanation for the difference in phenotypes in these mutant mice is that the *Kif6*^*Δ3*^ allele encodes a functional KIF6 protein. Analysis of these mutations in trans or quantitative analysis of these kinesin motor proteins *in vitro* is warranted to more fully address these conflicting observations.

There are marked differences in the phenotypes among the human, mouse, and zebrafish associated with mutations in *KIF6*. For example, *kif6* mutant zebrafish display post-natal onset scoliosis, mirroring adolescent idiopathic scoliosis (IS) in humans [29]. The formation of IS-like defects in zebrafish has been shown to be the result of a loss of CSF flow, associated with a loss of EC cilia and ventricular dilation during a defined window of larval zebrafish development [24]. Interestingly, we did not observe scoliosis in the *Kif6* mutant mice (S7 Fig), despite being of an appropriate age when IS-like scoliosis can manifest in mouse [30]. Moreover, we do not observe scoliosis in the patient at the age of 10 years, though it is possible that he may yet develop scoliosis during adolescence. The mechanism behind these differences may reflect distinctions in the functional input of the ventricular system for spine stability amongst teleosts and amniotes.

Furthermore, while we observe a clear role for KIF6 in maintaining the ventricular system in mouse and zebrafish, the patient does not have obvious hydrocephalus. However, his relative macrocephaly and slightly enlarged ventricles by MRI (Fig 1B-D) may suggest an element of what is commonly referred to as arrested hydrocephalus [31]. The contribution of EC cilia beating to bulk CSF flow might be species dependent. For instance, the majority of CSF flow in humans is thought to occur via the generation of a source-sink gradients; partly from the secretion of the choroid plexus and exchanges of the interstitial fluids, coupled with absorption at the arachnoid villi and lymphatics [32]. In contrast, localized or near-wall CSF flow [4], generated by polarized beating of EC cilia, are clearly important for the formation of hydrocephalus in rodents [8]; however, there have been limited examples EC cilia defects causing hydrocephalus in humans. Regardless there is growing evidence suggesting that EC cilia dependent CSF flow is crucial for the regulation of brain function and neurogenesis [4], and for adult neural stem cell proliferation [9]. It is possible that a specific loss of EC cilia in humans may only have minor effects on CSF bulk flow and ventricular homeostasis, while causing severe defects of neurogenesis leading to intellectual disability and other neurological disease. It will be important to determine if the loss of KIF6 function in adulthood will contribute to changes in neurodevelopment and behavior, cognitive decline, and ventricular homeostasis.

Finally, *KIF6* now joins five other kinesin genes, *KIF1C*, *KIF2A*, *KIF4A*, *KIF5C* and *KIF7* that were previously reported to be associated with neurological abnormalities in humans [33–36]. Here we suggest that *KIF6* has a uniquely specific function in the EC cilia in vertebrates, resulting in both cognitive impairment and macrocephaly in a child with a homozygous one-base pair deletion. Using a cell biological approach, we identified specific loss of EC cilia in *Kif6* mutant models in both mouse and zebrafish, suggesting a strong conservation of KIF6 function in ventricular system in vertebrates.

## Material and Methods

### Identification of Mutation in Patient 1

#### Whole Exome Sequencing (WES)

The patient’s genomic DNA of patient was extracted from peripheral blood leukocyte using AchivePure DNA Blood Kit (5 Prime Inc., Gaithersburg, MD). The sample was sent to Macrogen, Inc. (Seoul, Korea) for whole exome sequencing. The 4 ug of DNA sample was enriched by TruSeq Exome Enrichment Kit and was sequenced onto Hiseq 2000. The raw data per exome was mapped to the human reference genome hg19 using CASAVA v1.7. Variants calling were detected with SAMtools.

#### Homozygosity mapping

The sample was sent to Macrogen, Inc. (Seoul, Korea) for genotyping. The DNA sample was genotyped by HumanOmni 2.5-4v1 DNA BeadChip (Illumina) which contain 2,443,177 SNPs. The experiment was performed by the array protocol. PLINK was used to analyze for the homozygous regions.

#### Mutation analysis

We performed resequencing of *KIF6* pathogenic region in patient and patient’s family. Primers for the amplification of the candidate variant were designed using Primer 3 software (version 0.4.0). Primers KIF6-1193delT-F 5’-CAGCTTGAACATGGCTGAAA-3’ and KIF6-1193delT-R 5’-TTCTGTAAAGAGGTGGGAACAA-3’were used to amplify. The 20 ul of PCR reaction contained 50-100 ng of genomic DNA, 200 uM of each dNTP, 150 nM of each primer, 1. 5 mM MgCl_2_ and 0.5 unit of Taq DNA polymerase (Fermentas Inc., Glen Burnie, MD). The PCR condition was started with 95 °C for 5 min for pre-denaturation following with the 35 cycles of 94 °C for 30 sec, 55 °C for 30 sec and 72 °C for 30 sec. The product size of these primers is 276 bp. For sequencing, PCR products were treated with ExoSAP-IT (USP Corporation, Cleveland, OH), and sent for direct sequencing at Macrogen Inc. (Seoul, Korea). Bi-directional sequencing was done by using KIF6-1193delT F and R primers. Analyses were performed by Sequencher 4.2 (Gene Codes Corporation, Ann Arbor, MI).

#### PCR-RFLP

One hundred chromosomes and patient’s trio were genotyped by PCR-RFLP. Primer KIF6 *MfeI* F 5’-TGGCTTCACTATAAATTTCACTTTGTCAATG-3’ and mutagenic primer KIF6 mutagenic MfeI R 5’-TCCTGGTCTTCCAAAAAGGATGCAAT-3’were used to amplify KIF6 T-deletion. The 20 ul of PCR reaction contained 50-100 ng of genomic DNA, 200 uM of each dNTP, 150 nM of each primer, 1.5 mM MgCl_2_ and 0.5 unit of Taq DNA polymerase (Fermentas Inc., Glen Burnie, MD). The PCR condition was started with 95 C for 5 min for pre-denaturation following with the 35 cycles of 94 C for 30 sec, 60 C for 30 sec and 72 C for 30 sec. The product size of these primers is around 223 bp. The PCR product was incubated with 10U of Mfe-HF (New England Biolabs, Ipswich, MA) at 37 C overnight. Three percent of agarose gel electrophoresis was used to detect the different cut sizes of PCR product. A 196 bp and 26 bp bands were present in one base deletion sample.

#### Mice

All mouse studies and procedures were approved by the Animal Studies Committee at the University of Texas at Austin (AUP-2015-00185). The *Kif6*^*p.G555fs*^ mutant mouse were developed using CRISPR-Cas9-mediated genome editing. Using the CHOP-CHOP online tool [37], we identified a suitable 20-nucleotide site (GGAGATGTCACTGGGACGCC) targeting exon 14 of mouse *Kif6* (ENSMUST00000162854.1) in order to generate a C-terminal truncation allele. The gene specific and universal tracrRNA oligonucleotides (S3 Table) were annealed, filled in with CloneAmp HiFi PCR premix, column purified, and directly used for *in vitro* transcription of single-guide RNAs (sgRNAs) with a T7 Polymerase mix (M0255A NEB). All sgRNA reactions were treated with RNAse free-DNAse. We utilized a ssDNA oligo (S3 Table) to insert a frameshift mutation in all three reading frames, along with 8-cutter restriction sites for genotyping (3-stop donor) [38] (Fig S2). The *Kif6* ex14 3-stop donor and mKif6-R2-ex14-T7 sgRNA were submitted for pronuclear injection at the University of Texas at Austin Mouse Genetic Engineering Facility (UT-MGEF) using standard protocols (https://www.biomedsupport.utexas.edu/transgenics). We confirmed segregation of the *Kif6*^*p.G555fs*^ allele using several methods including increased mobility on a high percentage electrophoresis gel, donor-specific primer PCR, or PmeI (NEB) digestion of the *Kif6* exon14 amplicon (S2 Fig and S3 Table). PCR products in isolated alleles were cloned to pCRII TOPO (ThermoFisher) to identify scarless integration of the 3-stop donor at the *Kif6* locus using gene specific flanking primers (S3 Table).

*Kif6-LacZ*^*tm1b*^ mice were generated by injection of embryonic stem cell clones obtained from the Knockout Mouse Project (KOMP) Repository. Three *Kif6^tm1a(KOMP)Mbp^* embryonic stem (ES) cell clones (KOMP: EPD0736_3_G01; EPD0736_3_H02; and EPD0736_3_A03) all targeting exon 4 of the *Kif6* gene with a promoter-driven targeting cassette for the generation of a ’Knockout-first allele’ [39]. Pronuclear injections of all clones were done using standard procedures established by the UT MGEF. After screening for germline transmission, we isolated and confirmed a single heterozygous founder male (*Kif6*^*tm1a(KOMP)Mbp*^) carrier derived from the G01 clone. We confirmed the locus by long-range PCR, several confirmation PCR strategies targeting specific transgene sequences, and Sanger sequencing of the predicted breakpoints (S3 Table). After several backcrosses to the WT C57BL/6J substrain (JAX), we crossed a hemizygous *Kif6*^*tm1a/+*^ mutant male to a homozyogus *CMV-Cre* female (*B6.C-Tg(CMV-cre)1Cgn/J*) (JAX, 006054) to convert the *Kif6*^*tm1a*^ allele to a stable LacZ expressing *Kif6*^*tm1b*^ allele (*Kif6-LacZ*^*tm1b*^). Mutant F1 offspring from this cross were backcrossed to WT C57BL6/J mice and the F2 progeny were genotyped to confirm the *Kif6-LacZ*^*m1b*^ allele and the presence/absence of the CMV-Cre transgene. A single founder *Kif6-LacZ*^*tm1b*^ with the desired genotype (*Kif6-LacZ*^*tm1b*^ hemizygous, Cre transgene absent) was used to expand a colony for spatial expression analysis.

*Kif6*^*tm1c*^ conditional ready mice were generated by outcross of the *Kif6*^*tm1a(KOMP)Mbp*^ allele described above to a ubiquitously expressed Flippase strain (*129S4/SvJaeSor-Gt(ROSA)26Sor*^*tm1(FLP1)Dym*^*/J*) (JAX, 003946). F1 offspring were genotyped and sequenced at several breakpoints to ensure proper flip recombination and a single F1 founder was used to backcross to C57B6/J for propagation of the *Kif6*^*tm1c*^ strain. Analysis of recombination of the floxed *Kif6*^*tm1c*^ was performed by crossing homozygous *Kif6*^*tmlc/tmlc*^ to a compound heterozygous *CMV-Cre*; *Kif6*^*tm1c/+*^ mouse. Recombination of the exon 4 of Kif6 was confirmed by PCR-gel electrophoresis analysis (S3 Table).

#### LacZ Staining Protocol

Mice were perfused with LacZ fixative and post fixed for 2 hours at RT. Whole brains were then stained in X-gal solution overnight at 37°C followed by post-fixation in 4% PFA overnight at 4°C. The samples were then prepped for cryosectioning in 30% sucrose/OCT and sectioned. Sections were counter stained in Nuclear Fast Red stain (Sigma).

#### X-ray Analyses of Mice

Radiographs of the mouse skeleton were generated using a Kubtec DIGIMUS X-ray system (Kubtec, T0081B) with auto exposure under 25 kV.

#### Zebrafish Manipulations and Transgenesis

All zebrafish studies and procedures were approved by the Animal Studies Committee at the University of Texas at Austin (AUP-2015-00187). Adult zebrafish of the AB were maintained and bred as previously described [40]. Individual fish were used for analysis and compared to siblings and experimental control fish of similar size and age. Independent experiments were repeated using separate clutches of animals. Strains generated for this study: Tg(Foxj1a:GFP)^dp1^. Previously published strains: *kif6*^*sko*^ [19]. Transgenic lines were generated using the Tol2-system as described before [41].

#### Mouse and Zebrafish Perfusions and Embedding of Brain Tissues

Mice were humanely euthanized by extended CO_2_ exposure and transferred to chemical hood where the mouse was perfused with buffered saline followed by 4% PFA. Whole brains were placed in 4% PFA 4 hours at RT, then at 4° C overnight. Zebrafish were euthanized by exposure to lethal, extended dose of Tricane (8%) followed by decapitation. Zebrafish brains were extracted and fixed in 4% PFA at 4° C overnight. For paraffin embedding, the fixed brains were embedded and cut using standard paraffin embedding and sectioning protocols. Paraffin sections were stained with standard hematoxylin-eosin solution.

For frozen sections both mouse or zebrafish brains were fixed as above and then equilibrated to 30% or 35% sucrose, respectively at 4° C overnight. Whole brains were then placed in O.C.T. Compound (Tissue-Tek) and flash in cold ethanol bath. All blocks were stored at -80° Celsius until sectioning on a cryostat (Leica). All sections were dried at RT for ~2hrs. and stored at -80°C until use.

#### Immunofluorescence Protocol for Frozen Brain Sections

Sectioned tissues were warmed at room temperature for ~1 hour, then washed thrice in 1xPBS + 0. 1% Tween (PBST). Antigen retrieval was hot citrate buffer (pH6.8). Blocking was done in 10% Normal goat serum (Sigma) in 1xPBST. Primary antibodies (S100B at 1:1,000, ab52642, Abcam; CD133(Prominin-1), 134A, 1/500; Gamma Tubulin, sc-17787, Santa Cruz (C-11), 1/500; Anti-GFP, SC9996, Santa Cruz, 1:1,000) were diluted in 10% NGSS, 1xPBST and allowed to bind overnight at 4°C in a humidified chamber. Secondary fluorophores (Alexa Fluor 488(A-11034); 568(A10042); and 647(A32728), 1:1,000, ThermoFisher) were diluted in 10% NGS; 1xPBST were allowed to bind at RT for ~1hr. We used Prolong gold with DAPI (Cell Signaling Technologies, 8961) to seal coverslips prior to imaging.

#### Iodine-contrast μCT

Zebrafish specimens were fixed overnight in 10% buffered formalin, washed thrice in diH2O and stained ~48 hours in 25% Lugol’s solution/75% distilled water. Specimens were scanned by the High-resoultion X-ray CT Facility (http://www.ctlab.geo.utexas.edu/) on an Xradia at 100kV, 10W, 3.5s acquisition time, detector 11.5 mm, source -37 mm, XYZ [816, 10425, -841], camera bin 2, angles ±180, 1261 views, no filter, dithering, no sample drift correction. Reconstructed with center shift 5.5, beam hardening 0.15, theta -7, byte scaling [-150, 2200], binning 1, recon filter smooth (kernel size =0.5).

#### Statistical Analysis and image measures

GraphPad Prism version 7.0c for Mac (GraphPad Software) was used to analyze and plot data. Images for measurement were opened in FIJI (Image J) [42], and measures were taken using the freehand tool to draw outlines on ventricular area or whole brain area. Statistically significant differences between any two groups were examined using a two-tailed Student’s t-test, given equal variance. P values were considered significant at or below 0.05.

## ACKNOWLEDGMENTS

We thank members of the Gray and Wallingford labs for helpful discussions and critical reading of the manuscript. We would also like to thank Jin Xiang Ren and William Shawlot for help with mouse engineering, Terry Heckmann and Ryoko Minowa for excellent technical support. We thank Jessie Maisano for her help and expertise with iodine-contrast μCT for this study. This study was supported (V.S.) by the Thailand Research Fund (DPG6180001) and the Chulalongkorn Academic Advancement into Its 2nd Century Project. The research of M.K. was supported in part by the Provost Graduate Excellence Fellowship, Institute of Cell and Molecular Biology, University of Texas at Austin. The research of R.S.G. was supported start-up funds from the University of Texas at Austin Dell Medical School and by a NIH grant R01AR072009.

## Supporting Information

**S1 Fig.**
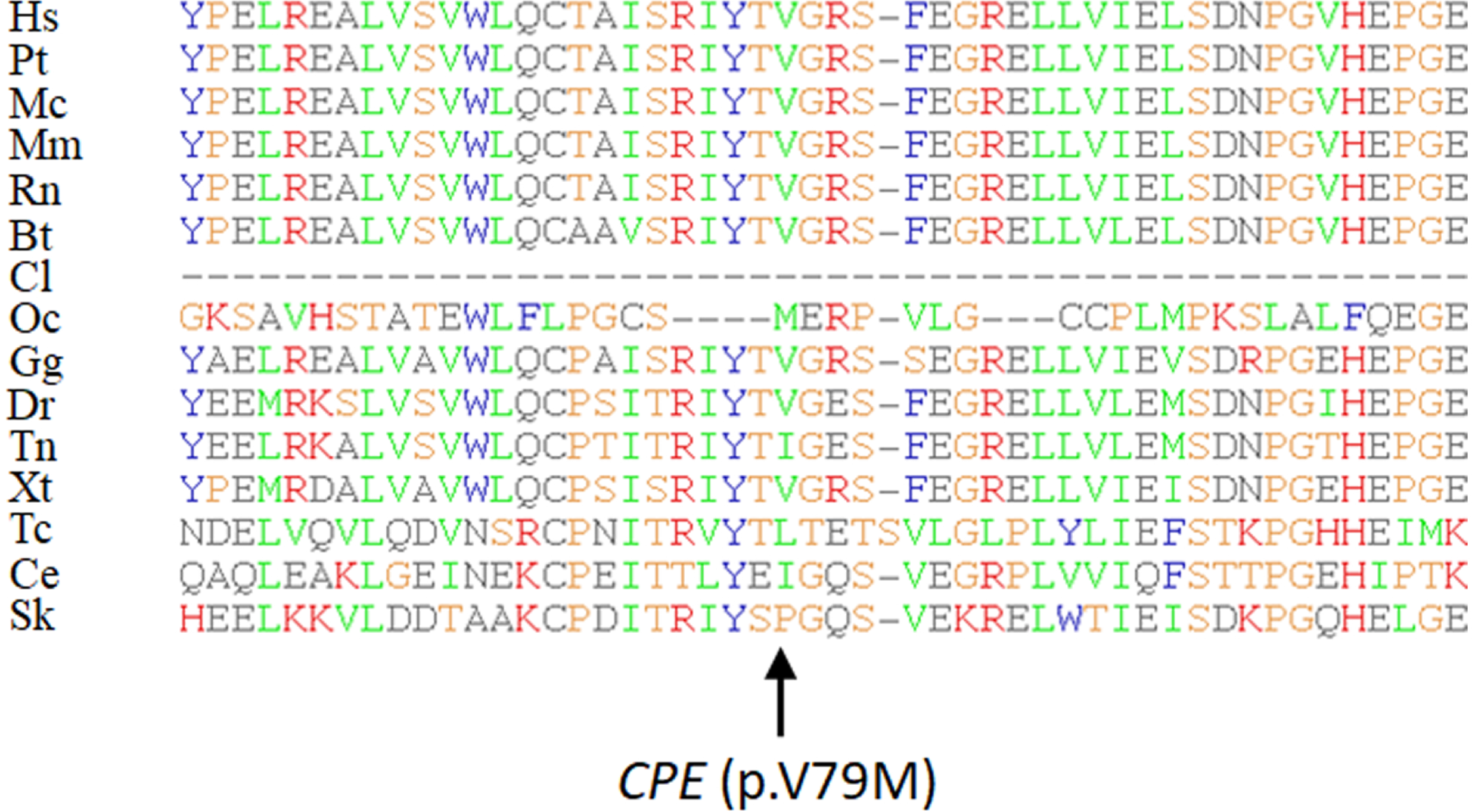
Clustal Alignment of *CPE* (p.V79M) variant. Hs, *Homo sapiens*; Pt, *Pan troglodytes*; Mc, *Macaca mulatta*; Mu, *Mus musculus*; Rn, *Rattus norvegicus*; Bt, *Bos Taurus*; Cl, *Canis lupus familiaris*; Oc, *Oryctolagus cuniculus*; Gg, *Gallus gallus*; Dr, *Danio rerio*; Tn, *Tetraodon nigroviridis*; Xt, *Xenopus (Silurana) tropicalis*; Tc, *Tribolium castaneum*; Ce, *Caenorhabditis elegans;* Sk, *Saccoglossus kowalevskii.*

**S2 Fig.**
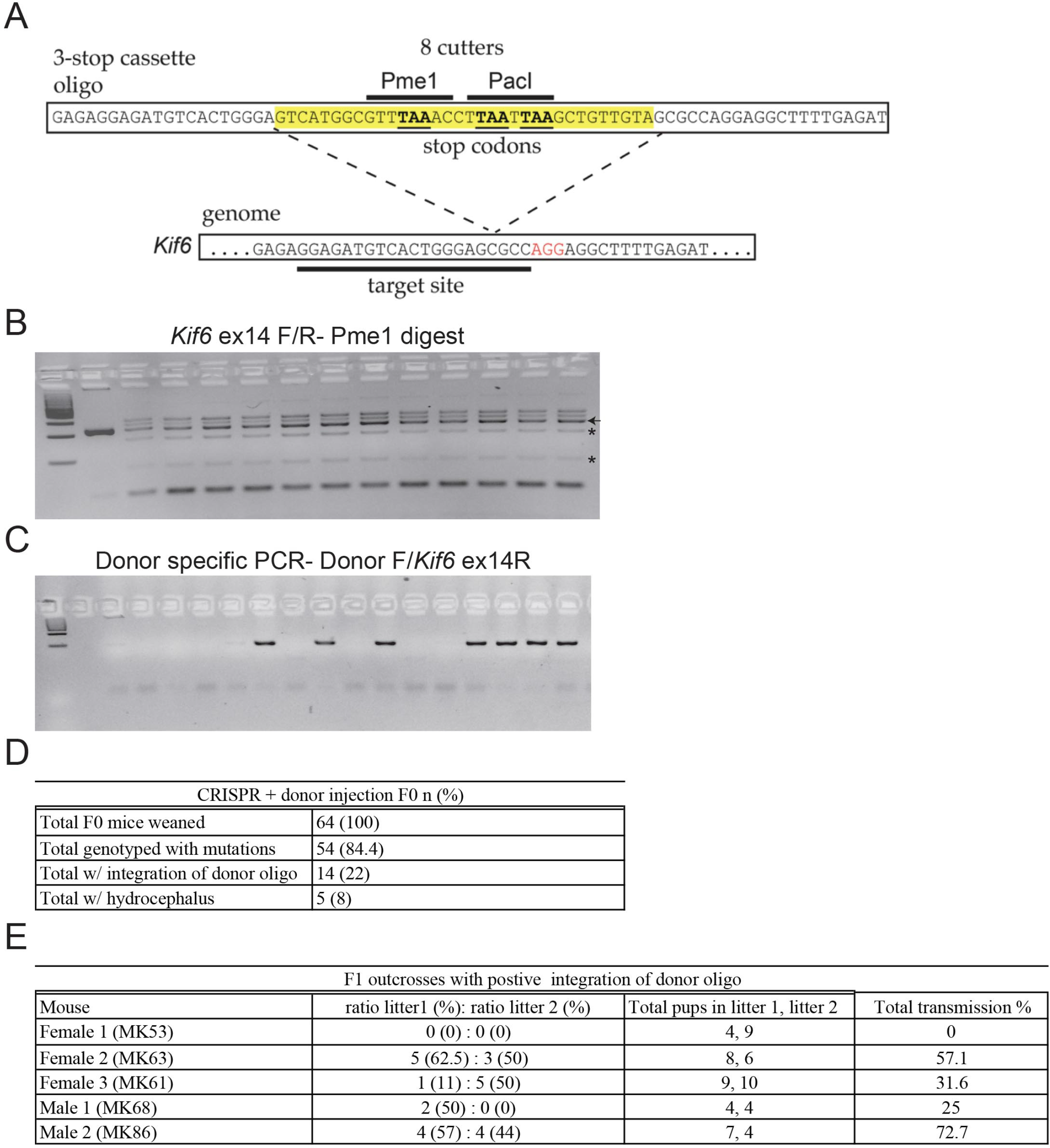
*Kif6*^*p.G555fs*^ generation and genotyping. (A) Schematic of target cut site and insertion cassette into exon 14 of *Kif6* locus. Insertion cassette contains three stop codons, one in each reading frame, and two 8 basepair restriction enzyme cut sites for easy genotyping. (B-C) Agarose gels of PCR products confirming germline transmission of donor cassette in F_1_ generation from CRISPR injected chimeras. (B) RE digest of PCR product from exon 14 flanking target site, shows cutting (asterisks) in heterozygous F1 mice. Wildtype band (arrow) appears in lane one and all the subsequent lanes. (C) PCR product from donor specific primer and *Kif6* exon 14 reverse primer confirming donor insertion and germline transmission. (D) Table describing CRISPR injected mice, number with detectable indels, total with integration of donor oligo, and total displaying hydrocephaly of chimeric injected CRISPR mice. (E) Germline transmission of donor cassette from chimeric CRISPR F0 mice to F1 generation.

**S3 Fig.**
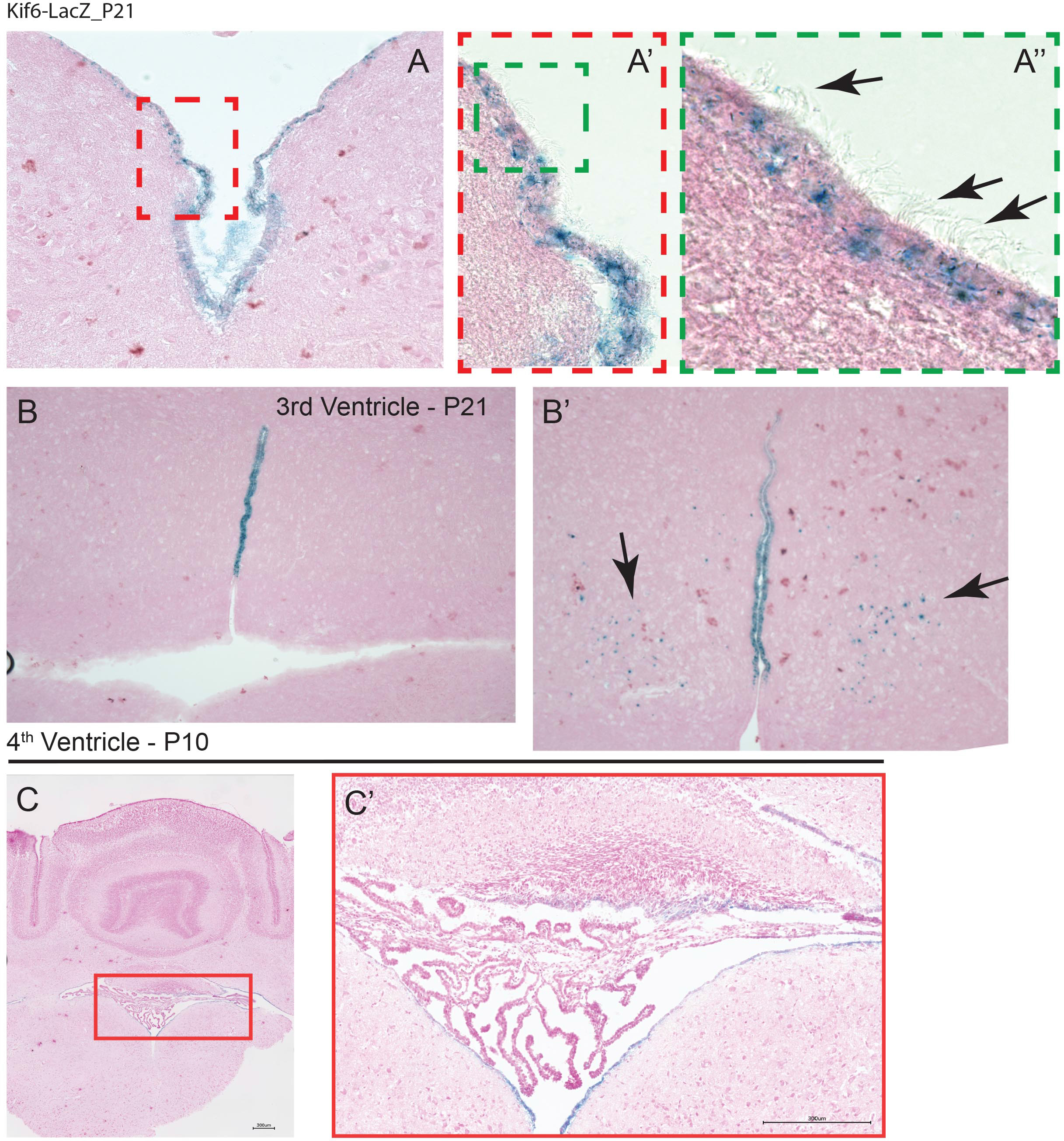
*LacZ* expression in different ages of *Kif6-LacZ*^*tmlb*^ transgenic mouse brain. (A-A’’) Coronal sections of P21 mouse brains showing LacZ staining restricted to the EP cell layer in the 4^th^ ventricle. Zoom in shows LacZ positive cells have cilia projecting into the lumen (arrows). (B-B’) Coronal sections of P21 mouse brains showing LacZ staining of ventral portion of 3^rd^ ventricle. (B’) Some sporadic staining appearing in the nuclei of the hypothalamus (arrows). (C-C’) LacZ staining in the fourth ventricle at P10 showing staining specific to ependymal cell layer.

**S4 Fig.**
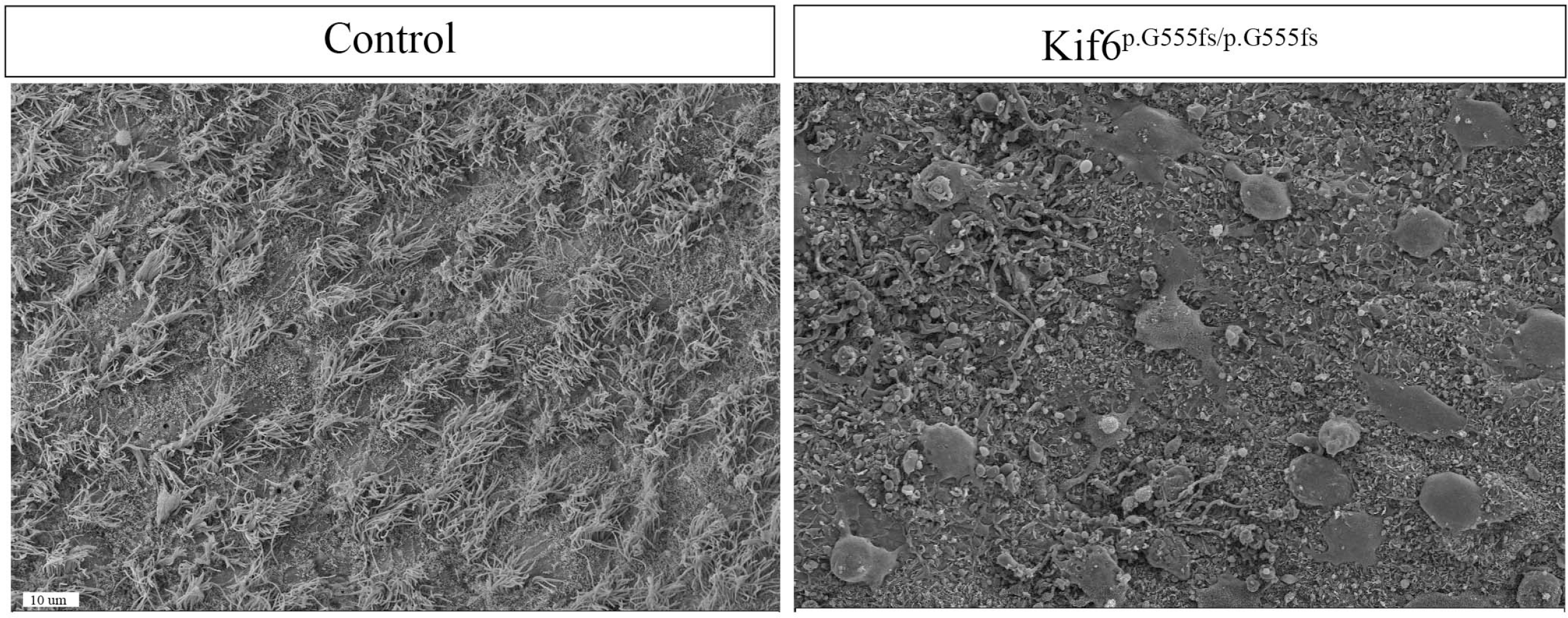
SEM of lateral ventricle in *Kif6*^*p.G555fs/p.G555fs*^ mutant and control at P28. SEM of *Kif6* wildtype vs. *Kif6*^*p.G555fs/p.G555fs*^ mutants shows *Kif6* mutants show a complete loss of ependymal cell cilia on the lateral wall by P28. Scale bar 10μM.

**S5 Fig.**
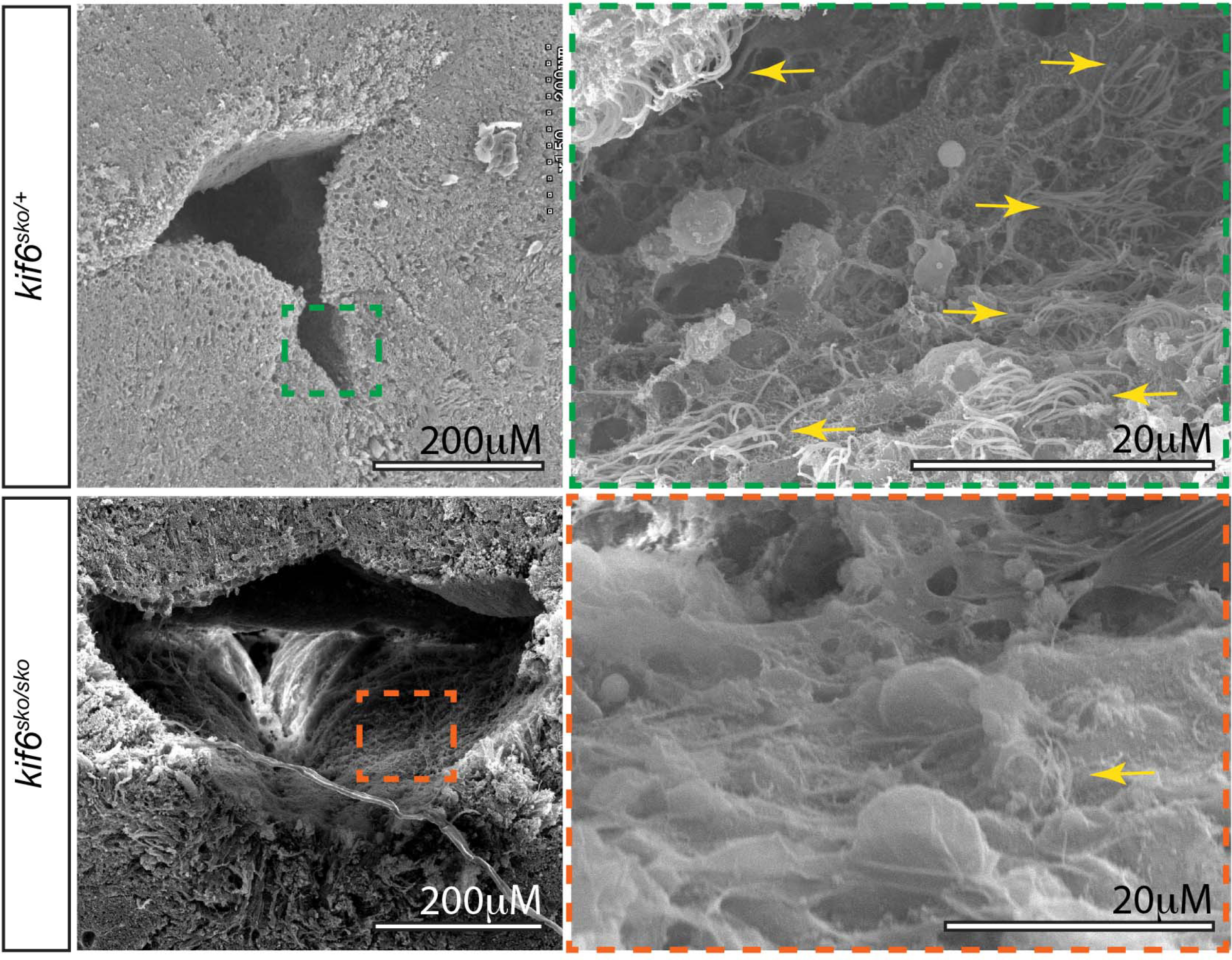
SEM of ventricle in *kif6*^*sko/sko*^ mutant zebrafish display dilation of the ventricular system and loss of ependymal cell cilia. SEM of zebrafish brain shows dilation of brain ventricles indicative of hydrocephaly. Higher magnification images reveal loss of ependymal cell cilia tufts in *kif6* zebrafish mutants when compared with heterozygous counterparts. Scale bars 20μM and 200μM.

**S6 Fig.**
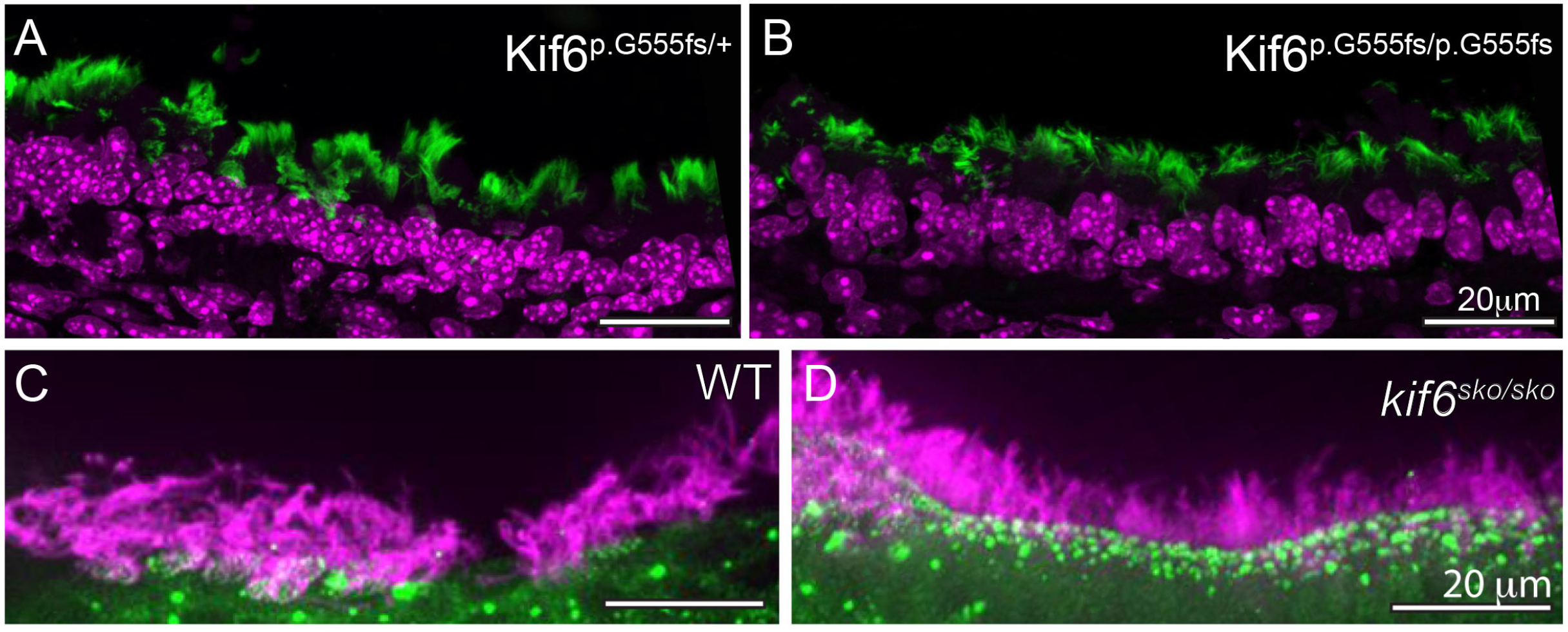
Immunofluoresence (IF) of Kif6 mutant multiciliated tissues in mouse and zebrafish. (A-B) Representative IF of trachea sections in *Kif6*^*p.G555fs/+*^ and *Kif6*^*p.G555fs/p.G555fs*^ mice showing no cilia defects present in trachea of *Kif6* mutant mice. Acetylated tubulin (green) marking cilia, DAPI-stained nuclei (magenta) (C-D) Representative IF of zebrafish nasal pit cilia shows typical cilia in *kif6* mutant zebrafish to wildtype counterparts. Acetylated tubulin (magenta) marking cilia, gamma-tubulin marking basal bodies (green). Scale bars are 20μM.

**S7 Fig.**
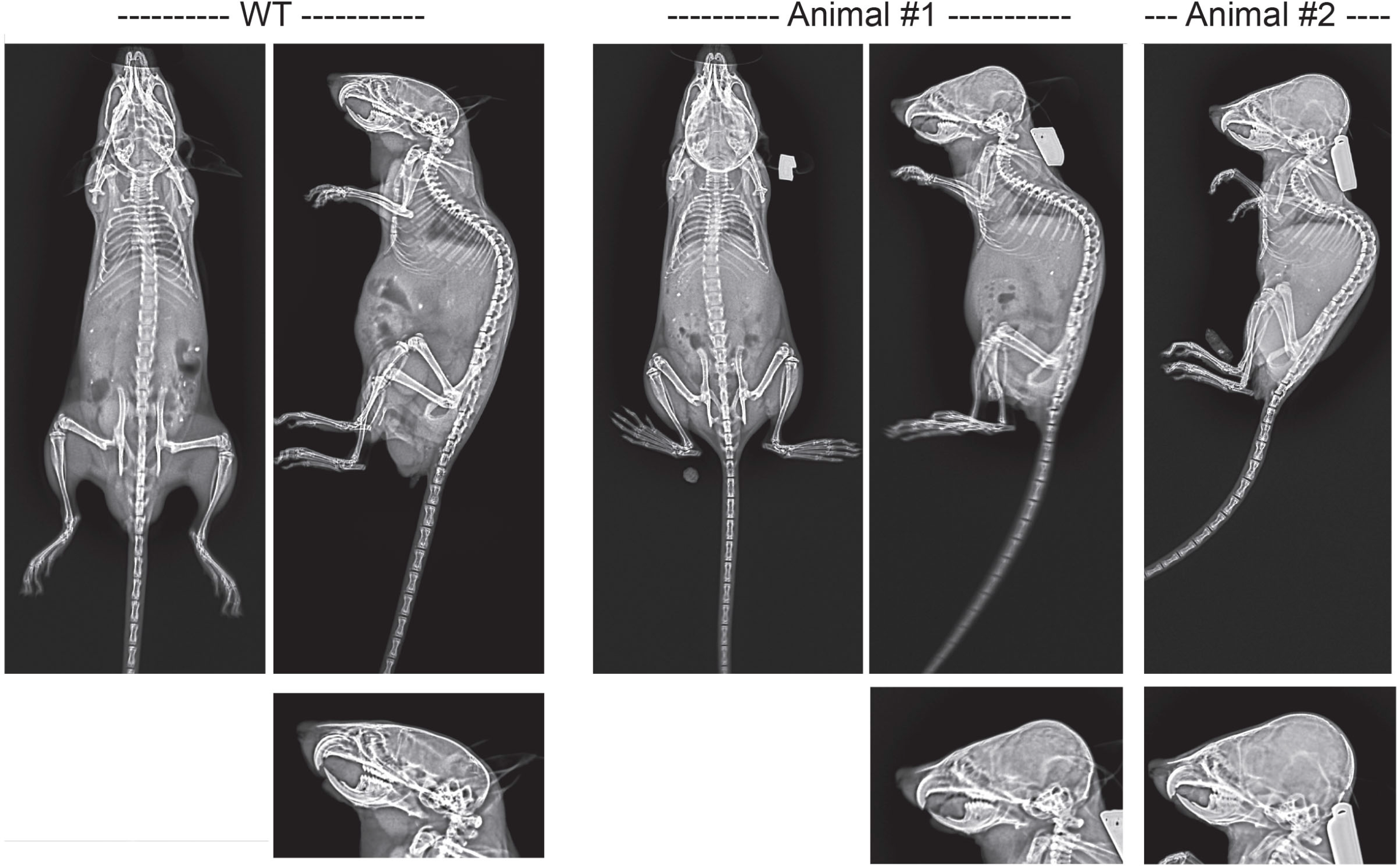
X-rays of *Kiftf*^*p.G555fs/p.G555fs*^ mutant mice. Representative X-rays of wildtype and *Kif6*^*p.G555fs/p.G555fs*^ mutant mice shows no scoliosis at P28. *Kif6*^*p.G555fs/p.G555fs*^ mice do however display skull expansion caused by progressive hydrocephalus (Animal #1 and #2).

**Table SI –.**
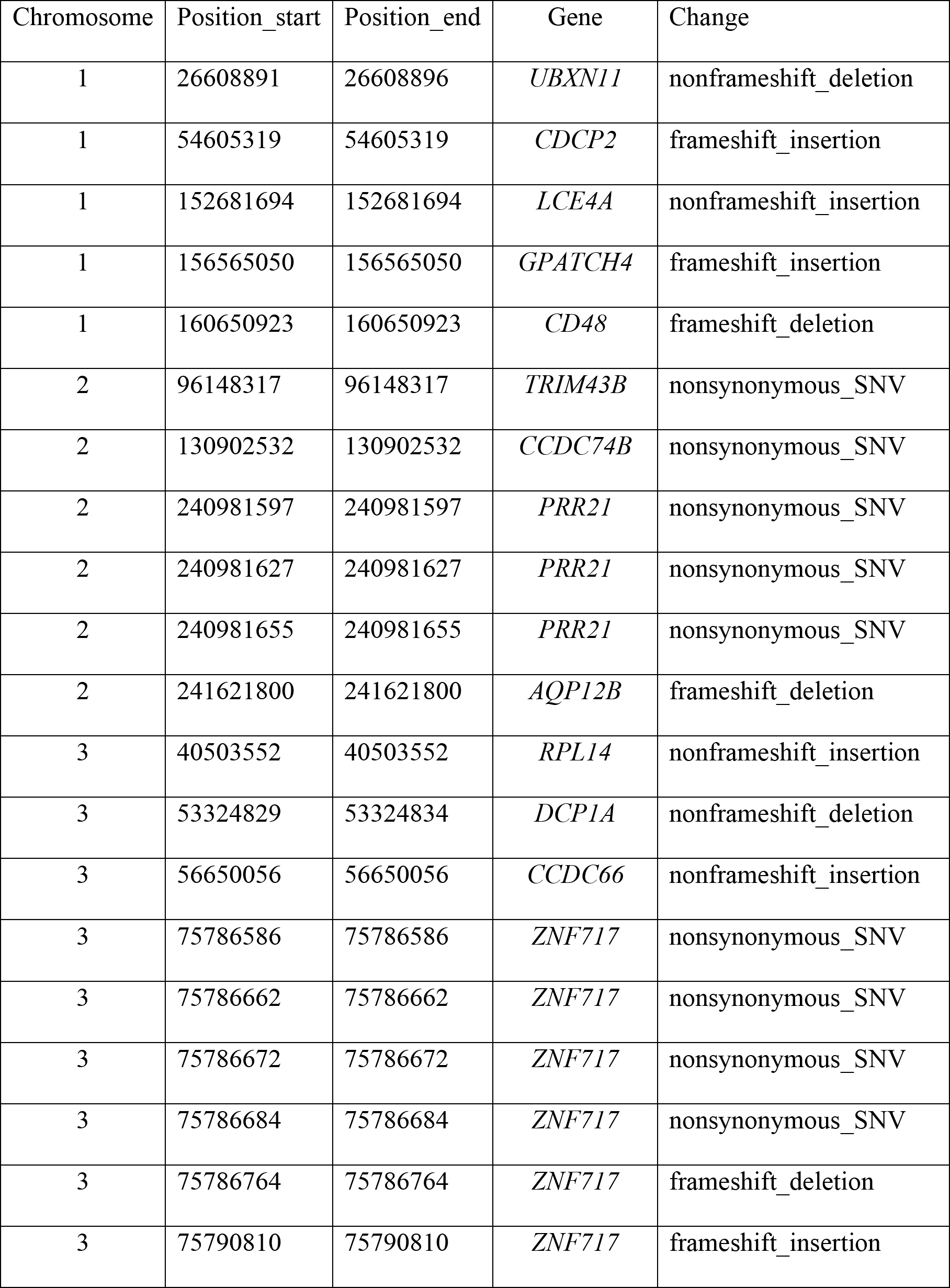
Eighty-three homozygous variants from WES.

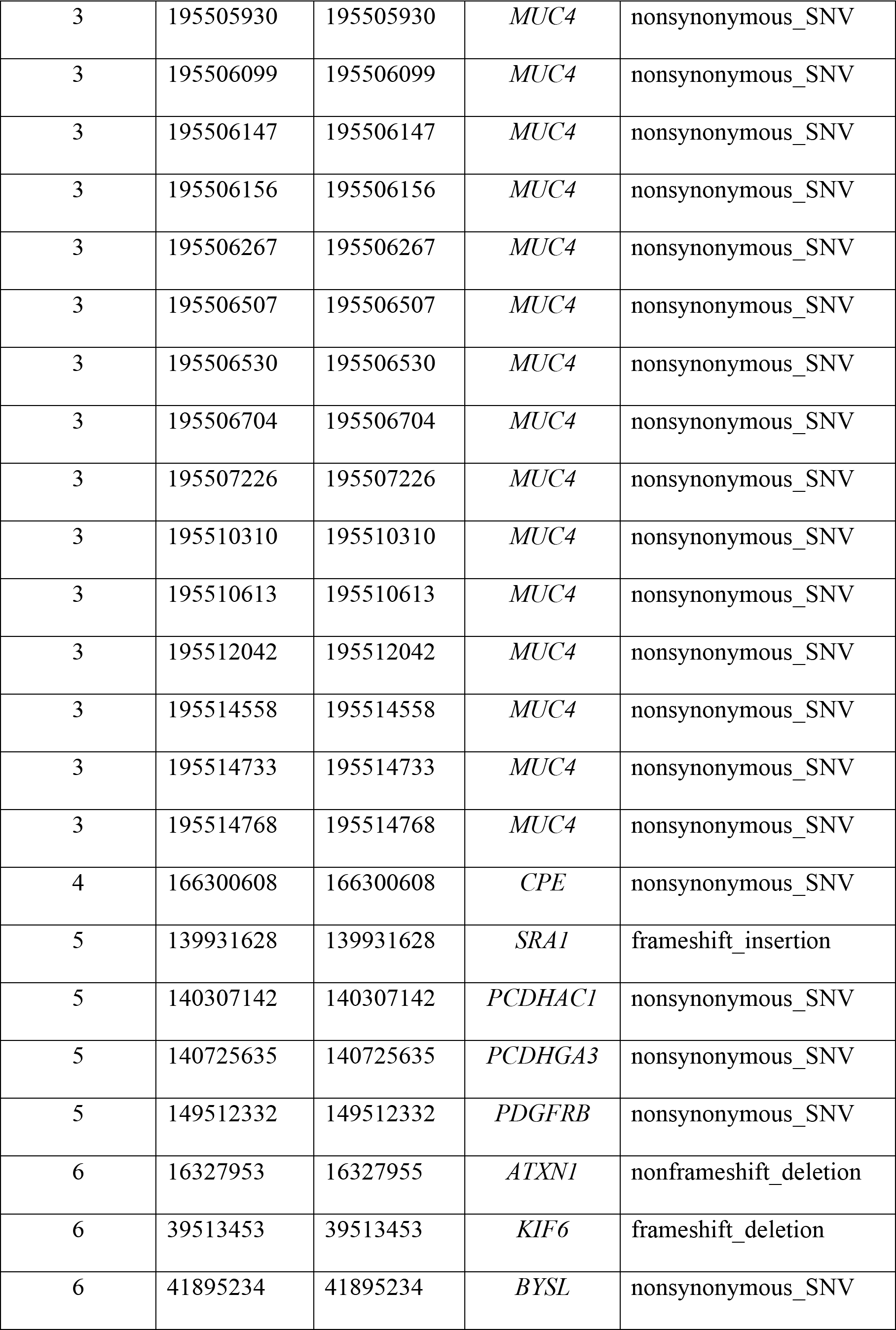

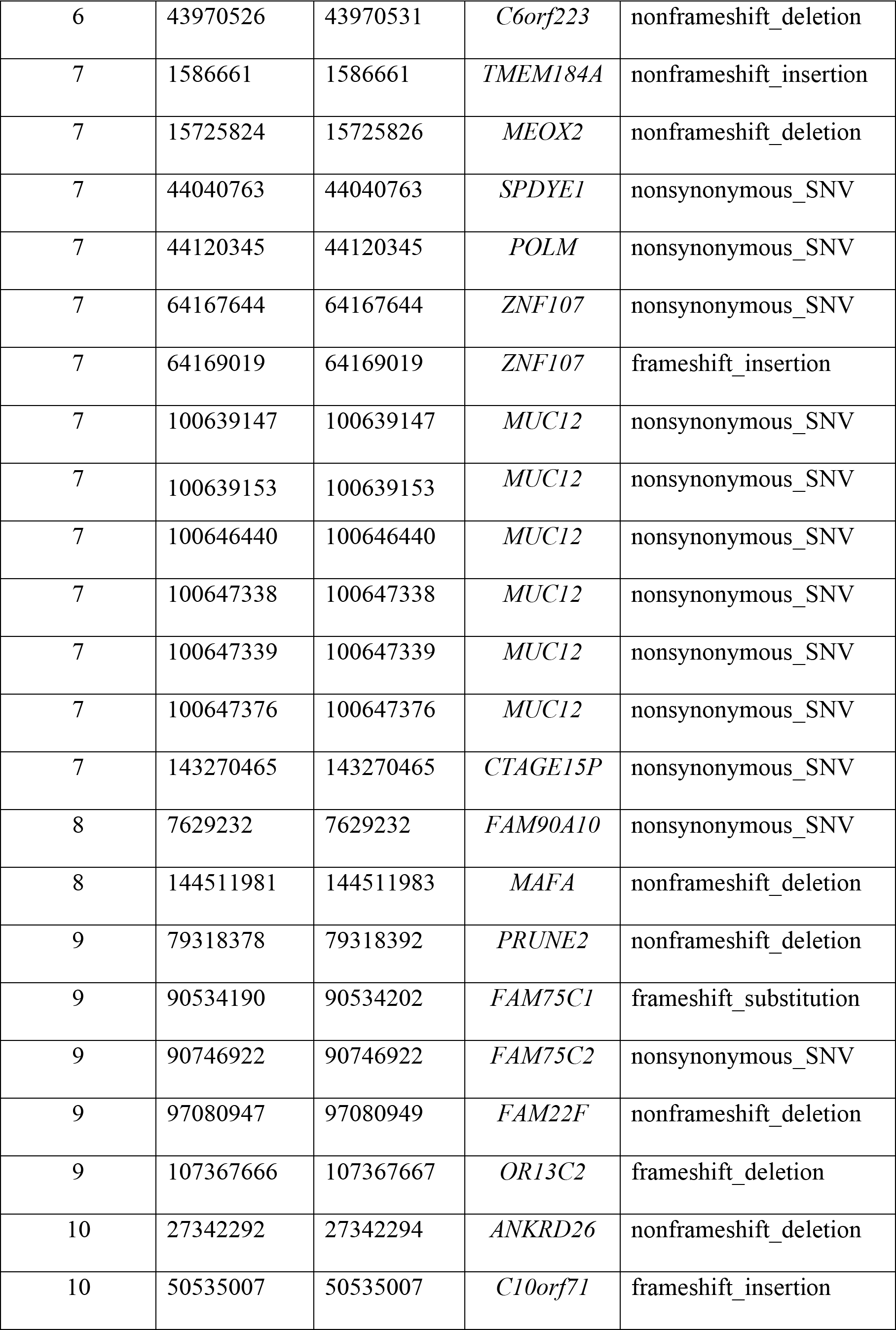

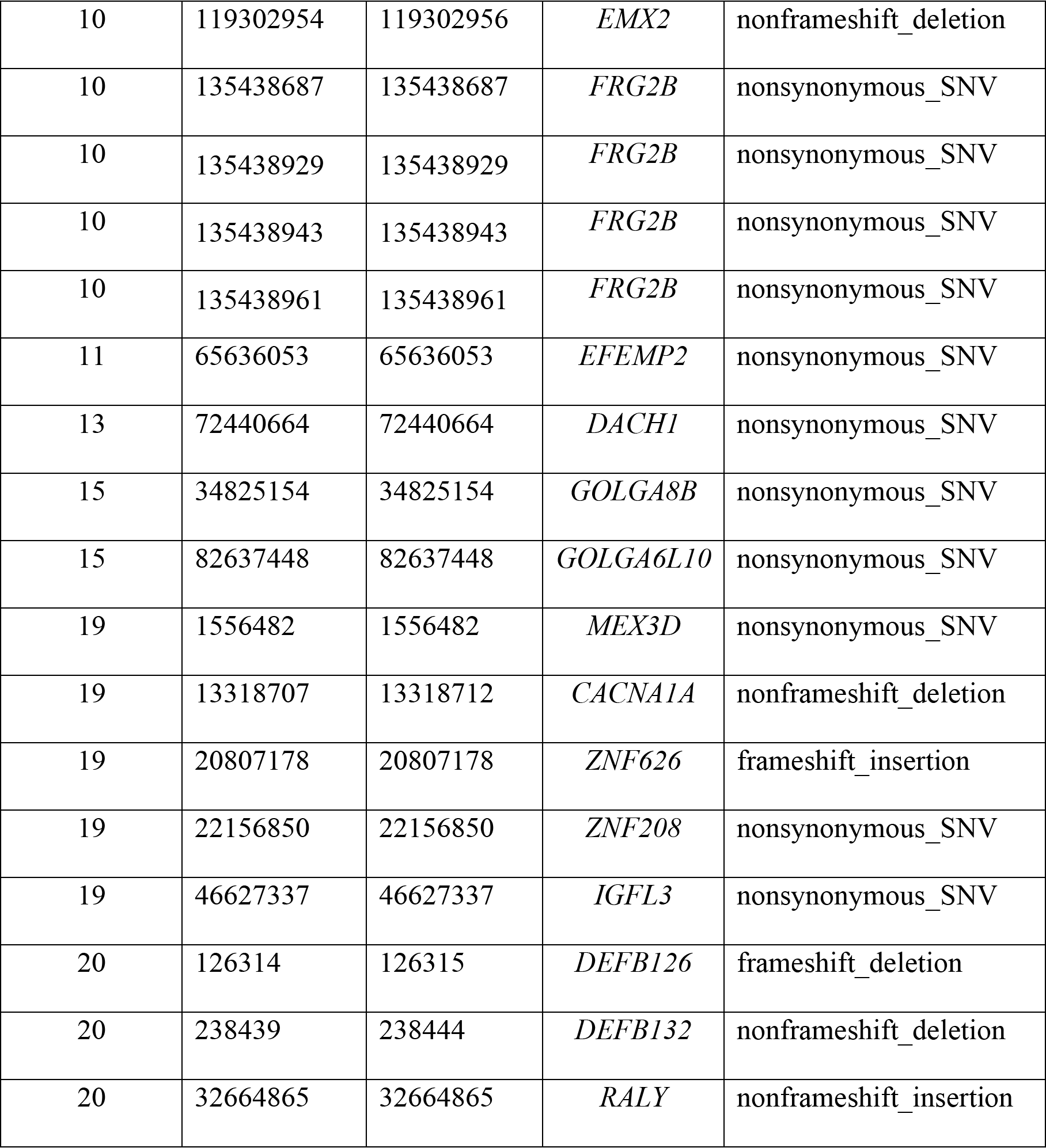

**Table SII –.**
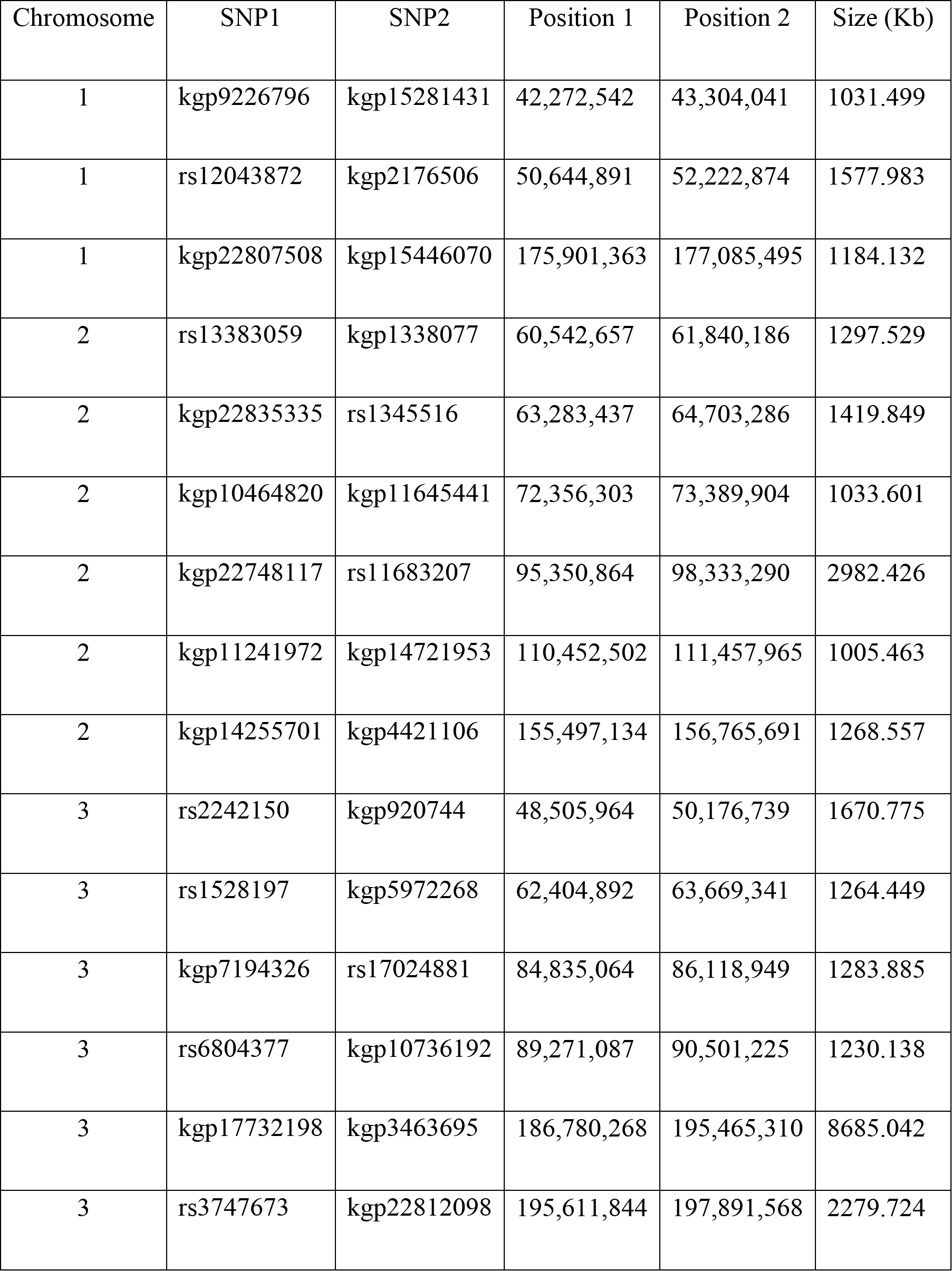
Sixty-three homozygous regions from homozygosity mapping.

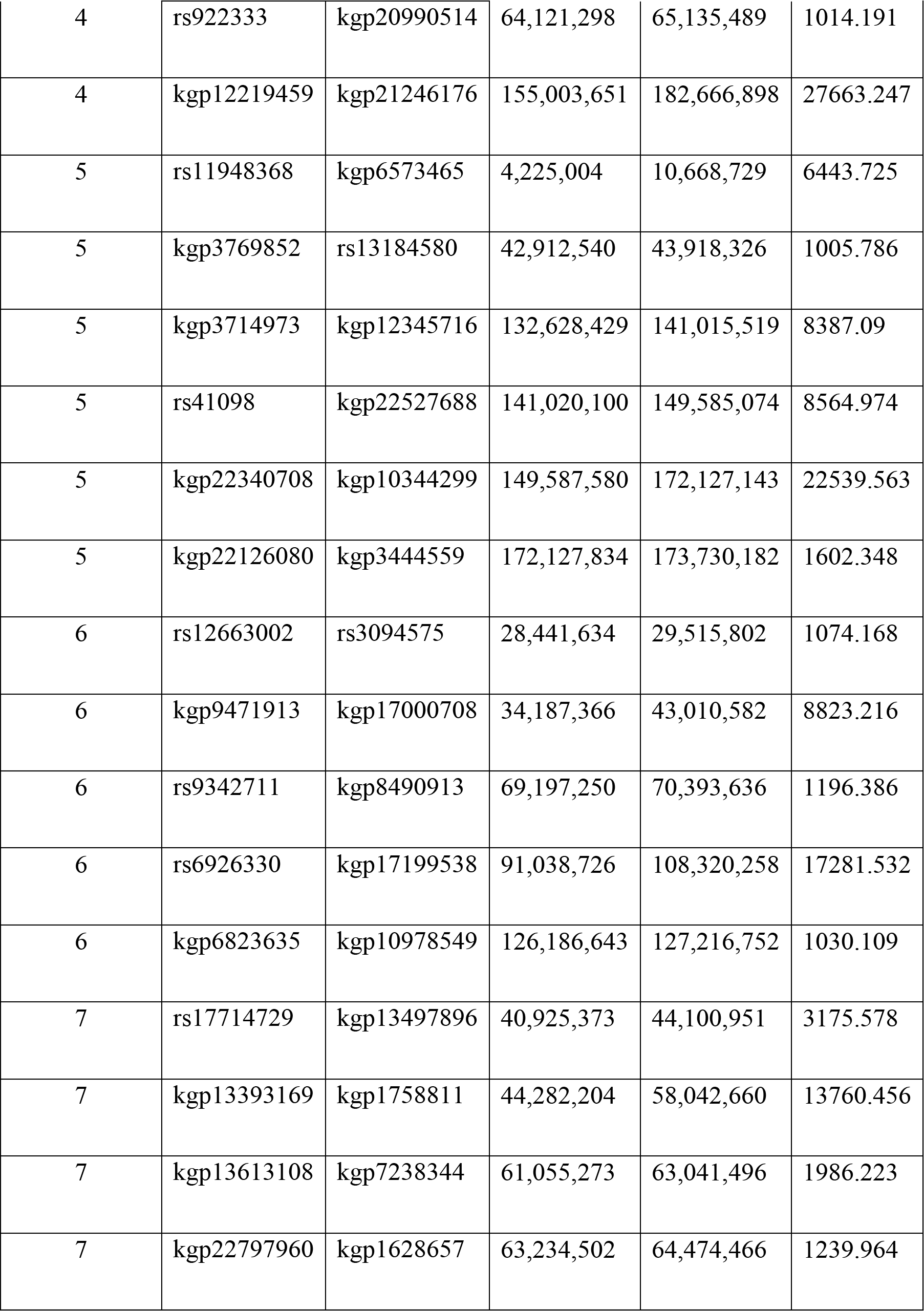

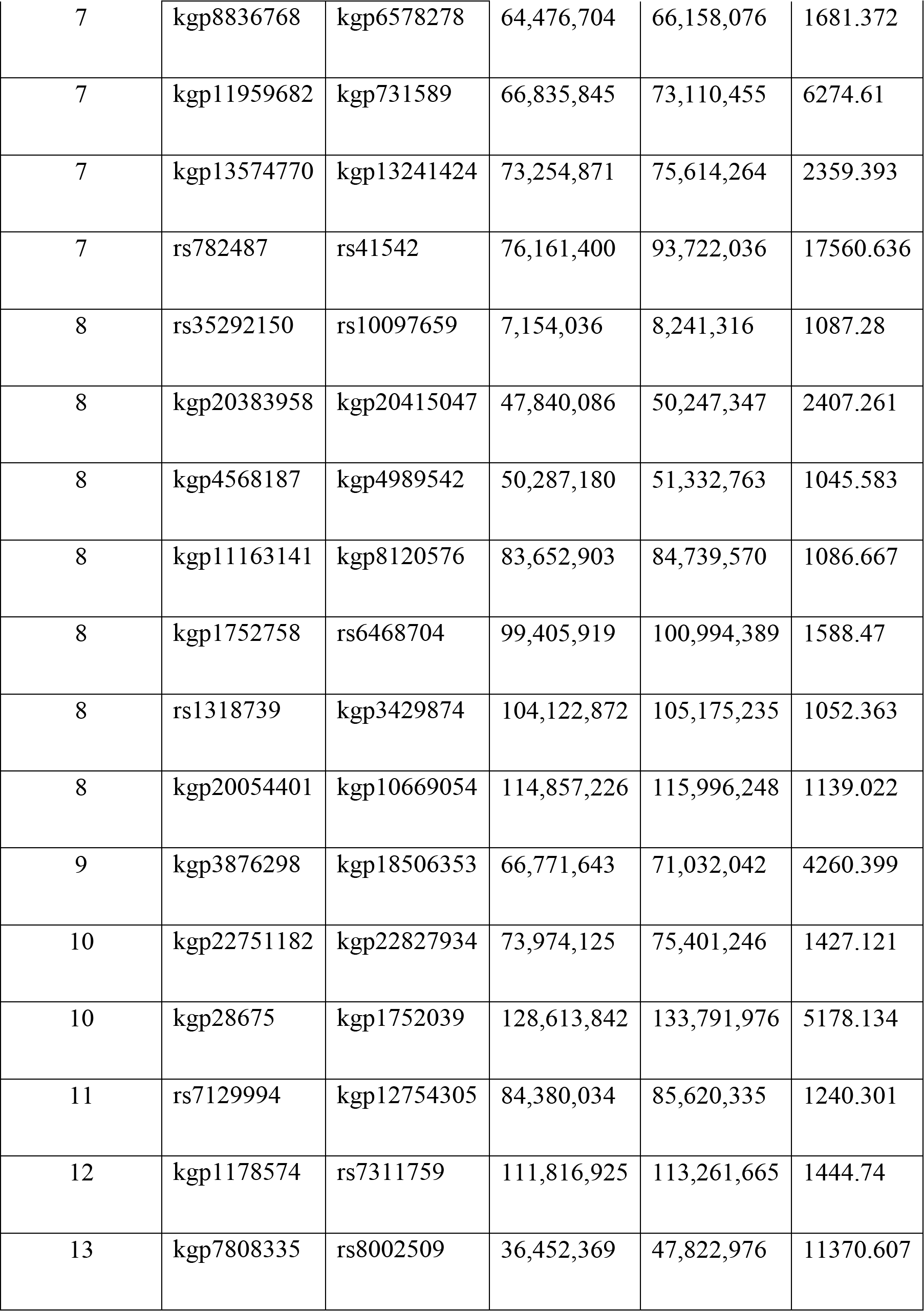

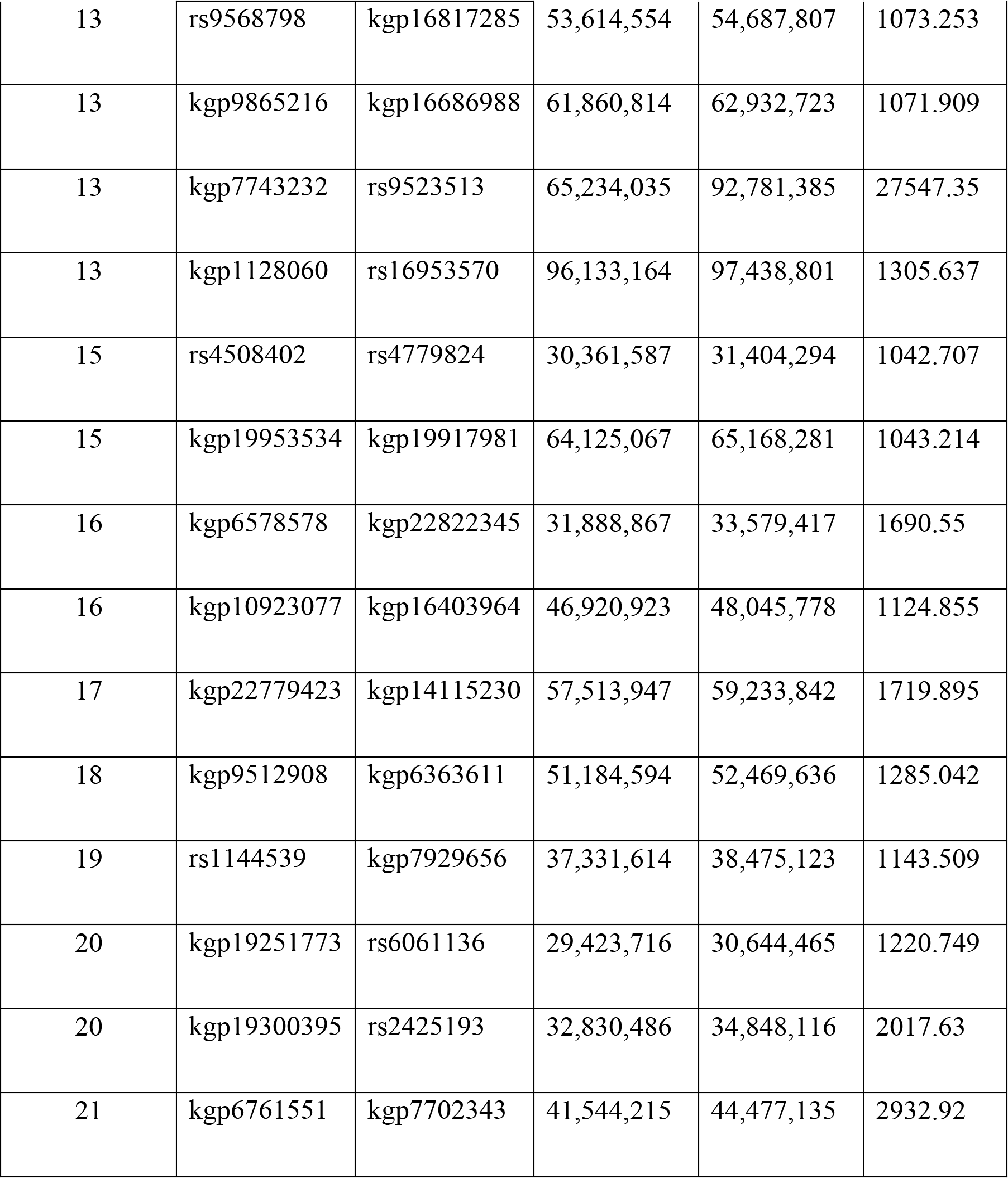

**Table SIII –.**
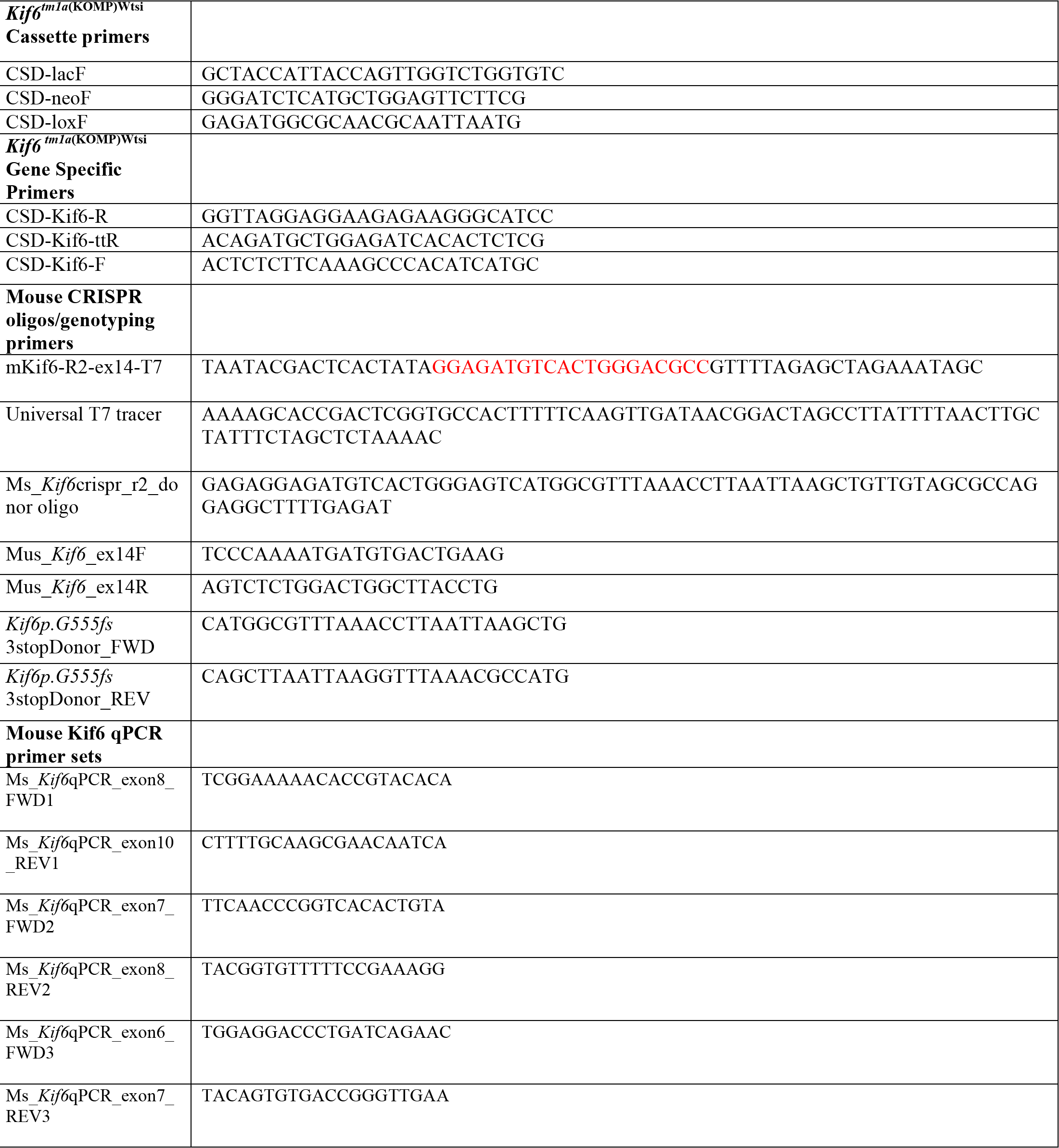
Mouse specific oligos and primers.

**Movie 1- Representative WT_Danio_Iodine-contrasted microCT transverse**

**Movie 2- Representative *kif6*^*sko/sko*^_Danio_Iodine-contrasted microCT transverse**

## References

1. Whedon JM, Glassey D. Cerebrospinal fluid stasis and its clinical significance. Altern Ther Health Med. 2009;15(3):54–60. PubMed PMID: 19472865; PubMed Central PMCID: PMCPMC2842089.

2. Rubenstein E. Relationship of senescence of cerebrospinal fluid circulatory system to dementias of the aged. Lancet. 1998;351(9098):283–5. doi:10.1016/S0140-6736(97)09234-9. PubMed PMID: 9457114.

3. Jacquet BV, Salinas-Mondragon R, Liang H, Therit B, Buie JD, Dykstra M, et al. FoxJ1-dependent gene expression is required for differentiation of radial glia into ependymal cells and a subset of astrocytes in the postnatal brain. Development. 2009;136(23):4021–31. doi:10.1242/dev.041129. PubMed PMID: 19906869; PubMed Central PMCID: PMCPMC3118431.

4. Spassky N, Meunier A. The development and functions of multiciliated epithelia. Nat Rev Mol Cell Biol. 2017;18(7):423–36. Epub 2017/04/13. doi:10.1038/nrm.2017.21. PubMed PMID: 28400610.

5. Gray RS, Roszko I, Solnica-Krezel L. Planar cell polarity: coordinating morphogenetic cell behaviors with embryonic polarity. Dev Cell. 2011;21(1):120–33. doi:10.1016/j.devcel.2011.06.011. PubMed PMID: 21763613; PubMed Central PMCID: PMCPMC3166557.

6. Ohata S, Nakatani J, Herranz-Perez V, Cheng J, Belinson H, Inubushi T, et al. Loss of Dishevelleds disrupts planar polarity in ependymal motile cilia and results in hydrocephalus. Neuron. 2014;83(3):558–71. doi:10.1016/j.neuron.2014.06.022. PubMed PMID: 25043421; PubMed Central PMCID: PMCPMC4126882.

7. Banizs B, Pike MM, Millican CL, Ferguson WB, Komlosi P, Sheetz J, et al. Dysfunctional cilia lead to altered ependyma and choroid plexus function, and result in the formation of hydrocephalus. Development. 2005;132(23):5329–39. doi:10.1242/dev.02153. PubMed PMID: 16284123.

8. Lee L. Riding the wave of ependymal cilia: genetic susceptibility to hydrocephalus in primary ciliary dyskinesia. J Neurosci Res. 2013;91(9):1117–32. doi:10.1002/jnr.23238. PubMed PMID: 23686703.

9. Petrik D, Myoga MH, Grade S, Gerkau NJ, Pusch M, Rose CR, et al. Epithelial Sodium Channel Regulates Adult Neural Stem Cell Proliferation in a Flow-Dependent Manner. Cell Stem Cell. 2018;22(6):865–78 e8. Epub 2018/05/22. doi:10.1016/j.stem.2018.04.016. PubMed PMID: 29779889.

10. Kousi M, Katsanis N. The Genetic Basis of Hydrocephalus. Annu Rev Neurosci. 2016;39:409–35. doi:10.1146/annurev-neuro-070815-014023. PubMed PMID: 27145913.

11. Reiter JF, Leroux MR. Genes and molecular pathways underpinning ciliopathies. Nat Rev Mol Cell Biol. 2017;18(9):533–47. Epub 2017/07/1. doi:10.1038/nrm.2017.60. PubMed PMID: 28698599; PubMed Central PMCID: PMCPMC5851292.

12. Verhey KJ, Kaul N, Soppina V. Kinesin assembly and movement in cells. Annu Rev Biophys. 2011;40:267–88. doi:10.1146/annurev-biophys-042910-155310. PubMed PMID: 21332353.

13. Hirokawa N, Noda Y, Tanaka Y, Niwa S. Kinesin superfamily motor proteins and intracellular transport. Nat Rev Mol Cell Biol. 2009;10(10):682–96. doi:10.1038/nrm2774. PubMed PMID: 19773780.

14. Lechtreck KF. IFT-Cargo Interactions and Protein Transport in Cilia. Trends Biochem Sci. 2015;40(12):765–78. Epub 2015/10/27. doi:10.1016/j.tibs.2015.09.003. PubMed PMID: 26498262; PubMed Central PMCID: PMCPMC4661101.

15. Demonchy R, Blisnick T, Deprez C, Toutirais G, Loussert C, Marande W, et al. Kinesin 9 family members perform separate functions in the trypanosome flagellum. J Cell Biol. 2009;187(5):615–22. Epub 2009/12/02. doi:10.1083/jcb.200903139. PubMed PMID: 19948486; PubMed Central PMCID: PMCPMC2806587.

16. Niwa S, Nakajima K, Miki H, Minato Y, Wang D, Hirokawa N. KIF19A is a microtubule-depolymerizing kinesin for ciliary length control. Dev Cell. 2012;23(6):1167–75. Epub 2012/11/22. doi:10.1016/j.devcel.2012.10.016. PubMed PMID: 23168168.

17. Li Y, Iakoubova OA, Shiffman D, Devlin JJ, Forrester JS, Superko HR. KIF6 polymorphism as a predictor of risk of coronary events and of clinical event reduction by statin therapy. Am J Cardiol. 2010;106(7):994–8. Epub 2010/09/22. doi:10.1016/j.amjcard.2010.05.033. PubMed PMID: 20854963.

18. Assimes TL, Holm H, Kathiresan S, Reilly MP, Thorleifsson G, Voight BF, et al. Lack of association between the Trp719Arg polymorphism in kinesin-like protein-6 and coronary artery disease in 19 case-control studies. J Am Coll Cardiol. 2010;56(19):1552–63. Epub 2010/10/12. doi:10.1016/j.jacc.2010.06.022. PubMed PMID: 20933357; PubMed Central PMCID: PMCPMC3084526.

19. Buchan JG, Gray RS, Gansner JM, Alvarado DM, Burgert L, Gitlin JD, et al. Kinesin family member 6 (kif6) is necessary for spine development in zebrafish. Dev Dyn. 2014;243(12):1646–57. doi:10.1002/dvdy.24208. PubMed PMID: 25283277.

20. Schultz J, Milpetz F, Bork P, Ponting CP. SMART, a simple modular architecture research tool: identification of signaling domains. Proc Natl Acad Sci U S A. 1998;95(11):5857–64. Epub 1998/05/30. PubMed PMID: 9600884; PubMed Central PMCID: PMCPMC34487.

21. Schwenk F, Baron U, Rajewsky K. A cre-transgenic mouse strain for the ubiquitous deletion of loxP-flanked gene segments including deletion in germ cells. Nucleic Acids Res. 1995;23(24):5080–1. Epub 1995/12/25. PubMed PMID: 8559668; PubMed Central PMCID: PMCPMC307516.

22. Spassky N, Merkle FT, Flames N, Tramontin AD, Garcia-Verdugo JM, Alvarez-Buylla A. Adult ependymal cells are postmitotic and are derived from radial glial cells during embryogenesis. J Neurosci. 2005;25(1):10–8. doi:10.1523/JNEUR0SCI.1108-04.2005. PubMed PMID: 15634762.

23. Pfenninger CV, Roschupkina T, Hertwig F, Kottwitz D, Englund E, Bengzon J, et al. CD133 is not present on neurogenic astrocytes in the adult subventricular zone, but on embryonic neural stem cells, ependymal cells, and glioblastoma cells. Cancer Res. 2007;67(12):5727–36. Epub 2007/06/19. doi:10.1158/0008-5472.CAN-07-0183. PubMed PMID: 17575139.

24. Grimes DT, Boswell CW, Morante NF, Henkelman RM, Burdine RD, Ciruna B. Zebrafish models of idiopathic scoliosis link cerebrospinal fluid flow defects to spine curvature. Science. 2016;352(6291):1341–4. doi:10.1126/science.aaf6419. PubMed PMID: 27284198.

25. Metscher BD. MicroCT for developmental biology: a versatile tool for high-contrast 3D imaging at histological resolutions. Dev Dyn. 2009;238(3):632–40. doi:10.1002/dvdy.21857. PubMed PMID: 19235724.

26. Wullimann MF, Rupp B, Reichert H. Neuroanatomy of the zebrafish brain: a topological atlas. Basel; Boston: Birkhäuser Verlag; 1996. vi, 144 p. p.

27. Kee HL, Dishinger JF, Blasius TL, Liu CJ, Margolis B, Verhey KJ. A size-exclusion permeability barrier and nucleoporins characterize a ciliary pore complex that regulates transport into cilia. Nat Cell Biol. 2012;14(4):431–7. doi:10.1038/ncb2450. PubMed PMID: 22388888; PubMed Central PMCID: PMCPMC3319646.

28. Hameed A, Bennett E, Ciani B, Hoebers LP, Milner R, Lawrie A, et al. No evidence for cardiac dysfunction in Kif6 mutant mice. PLoS One. 2013;8(1):e54636. doi:10.1371/journal.pone.0054636. PubMed PMID: 23355886; PubMed Central PMCID: PMCPMC3552957.

29. Liu Z, Gray RS. Animal models of idiopathic scoliosis. In: Kusumi K, Dunwoodie SL, editors. The Genetics and Development of Scoliosis. New York, NY: Springer Nature: Cham, Switzerland.; 2018.

30. Karner CM, Long F, Solnica-Krezel L, Monk KR, Gray RS. Gpr126/Adgrg6 deletion in cartilage models idiopathic scoliosis and pectus excavatum in mice. Hum Mol Genet. 2015;24(15):4365–73. doi:10.1093/hmg/ddv170. PubMed PMID: 25954032; PubMed Central PMCID: PMCPMC4492399.

31. Schick RW, Matson DD. What is arrested hydrocephalus? J Pediatr. 1961;58:791–9. Epub 1961/06/01. PubMed PMID: 13747587.

32. Brinker T, Stopa E, Morrison J, Klinge P. A new look at cerebrospinal fluid circulation. Fluids Barriers CNS. 2014;11:10. Epub 2014/05/13. doi:10.1186/2045-8118-11-10. PubMed PMID: 24817998; PubMed Central PMCID: PMCPMC4016637.

33. de Ligt J, Willemsen MH, van Bon BW, Kleefstra T, Yntema HG, Kroes T, et al. Diagnostic exome sequencing in persons with severe intellectual disability. N Engl J Med. 2012;367(20):1921–9. Epub 2012/10/05. doi:10.1056/NEJMoa1206524. PubMed PMID: 23033978.

34. Najmabadi H, Hu H, Garshasbi M, Zemojtel T, Abedini SS, Chen W, et al. Deep sequencing reveals 50 novel genes for recessive cognitive disorders. Nature. 2011;478(7367):57–63. Epub 2011/09/23. doi:10.1038/nature10423. PubMed PMID: 21937992.

35. Poirier K, Lebrun N, Broix L, Tian G, Saillour Y, Boscheron C, et al. Mutations in TUBG1, DYNC1H1, KIF5C and KIF2A cause malformations of cortical development and microcephaly. Nat Genet. 2013;45(6):639–47. Epub 2013/04/23. doi:10.1038/ng.2613. PubMed PMID: 23603762; PubMed Central PMCID: PMCPMC3826256.

36. Willemsen MH, Ba W, Wissink-Lindhout WM, de Brouwer AP, Haas SA, Bienek M, et al. Involvement of the kinesin family members KIF4A and KIF5C in intellectual disability and synaptic function. J Med Genet. 2014;51(7):487–94. Epub 2014/05/09. doi:10.1136/jmedgenet-2013-102182. PubMed PMID: 24812067.

37. Labun K, Montague TG, Gagnon JA, Thyme SB, Valen E. CHOPCHOP v2: a web tool for the next generation of CRISPR genome engineering. Nucleic Acids Res. 2016;44(W1):W272–6. doi:10.1093/nar/gkw398. PubMed PMID: 27185894; PubMed Central PMCID: PMCPMC4987937.

38. Gagnon JA, Valen E, Thyme SB, Huang P, Ahkmetova L, Pauli A, et al. Efficient mutagenesis by Cas9 protein-mediated oligonucleotide insertion and large-scale assessment of single-guide RNAs. PLoS One. 2014;9(5):e98186. doi:10.1371/journal.pone.0098186. PubMed PMID: 24873830; PubMed Central PMCID: PMC4038517.

39. Skarnes WC, Rosen B, West AP, Koutsourakis M, Bushell W, Iyer V, et al. A conditional knockout resource for the genome-wide study of mouse gene function. Nature. 2011;474(7351):337–42. doi:10.1038/nature10163. PubMed PMID: 21677750; PubMed Central PMCID: PMCPMC3572410.

40. Gray RS, Wilm TP, Smith J, Bagnat M, Dale RM, Topczewski J, et al. Loss of col8a1a function during zebrafish embryogenesis results in congenital vertebral malformations. Dev Biol. 2014;386(1):72–85. doi:10.1016/j.ydbio.2013.11.028. PubMed PMID: 24333517; PubMed Central PMCID: PMC3938106.

41. Kawakami K. Tol2: a versatile gene transfer vector in vertebrates. Genome Biol. 2007;8 Suppl 1:S7. doi:10.1186/gb-2007-8-s1-s7. PubMed PMID: 18047699; PubMed Central PMCID: PMCPMC2106836.

42. Schindelin J, Arganda-Carreras I, Frise E, Kaynig V, Longair M, Pietzsch T, et al. Fiji: an open-source platform for biological-image analysis. Nat Methods. 2012;9(7):676–82. doi:10.1038/nmeth.2019. PubMed PMID: 22743772; PubMed Central PMCID: PMCPMC3855844.

